# In-cell chemical crosslinking identifies hotspots for p62-IκBα interaction that underscore a critical role of p62 in limiting NF-κB activation through IκBα-stabilization

**DOI:** 10.1101/2022.10.13.512146

**Authors:** Yi Liu, Michael J. Trnka, Liang He, A. L. Burlingame, Maria Almira Correia

**Affiliations:** Departments of Cellular & Molecular Pharmacology, University of California San Francisco, San Francisco CA 94158-2517; Pharmaceutical Chemistry, University of California San Francisco, San Francisco CA 94158-2517; Bioengineering and Therapeutic Sciences, University of California San Francisco, San Francisco CA 94158-2517; The Liver Center, University of California San Francisco, San Francisco CA 94158-2517

**Keywords:** IκBα, NF-κB, p62/SQSTM-1, 20S/26S proteasomal degradation, chemical crosslinking mass spectrometry, APEX-proximity biotinylation labeling, liver inflammation, IL-1β, TNFα

## Abstract

We have previously documented that in liver cells, the multifunctional protein scaffold p62/SQSTM1 is closely associated with IκBα, an inhibitor of the transcriptional activator NF-κB. Such an intimate p62-IκBα association we now document leads to a marked 18-fold proteolytic IκBα-stabilization, enabling its nuclear entry and termination of the NF-κB-activation cycle. In p62^-/-^-cells, such termination is abrogated resulting in the nuclear persistence and prolonged activation of NF-κB following inflammatory stimuli. Utilizing various approaches both classic (structural deletion, site-directed mutagenesis) as well as novel (in cell chemical crosslinking), coupled with proteomic analyses, we have defined the precise structural hotspots of p62-IκBα association. Accordingly, we have identified such IκBα hotspots to reside around N-terminal (K38, K47 and K67) and C-terminal (K238/C239) residues in its 5^th^ ankyrin repeat domain. These sites interact with two hotspots in p62: One in its PB-1 subdomain around K13, and the other comprised of a positively charged patch (R_183_/R_186_/K_187_/K_189_) in the intervening region between its ZZ- and TB-subdomains. APEX proximity analyses upon IκBα co-transfection of cells with and without p62 have enabled the characterization of the p62 influence on IκBα-protein-protein interactions. Interestingly, consistent with p62’s capacity to proteolytically stabilize IκBα, its presence greatly impaired IκBα’s interactions with various 20S/26S proteasomal subunits. Furthermore, consistent with p62-interaction with IκBα on an interface opposite to that of its NF-κB-interacting interface, p62 failed to significantly affect IκBα-NF-κB interactions. These collective findings together with the known dynamic p62 nucleocytoplasmic shuttling, leads us to speculate that it may be involved in “piggy-back” nuclear transport of IκBα following its NF-κB-elicited transcriptional activation and *de novo* synthesis, required for the termination of the NF-κB-activation cycle. Consequently, mice carrying a liver specific deletion of p62-residues 68-252 harboring its positively charged patch, reveal age-dependent enhanced liver inflammation. Our findings reveal yet another mode of p62-mediated pathophysiologically relevant regulation of NF-κB.

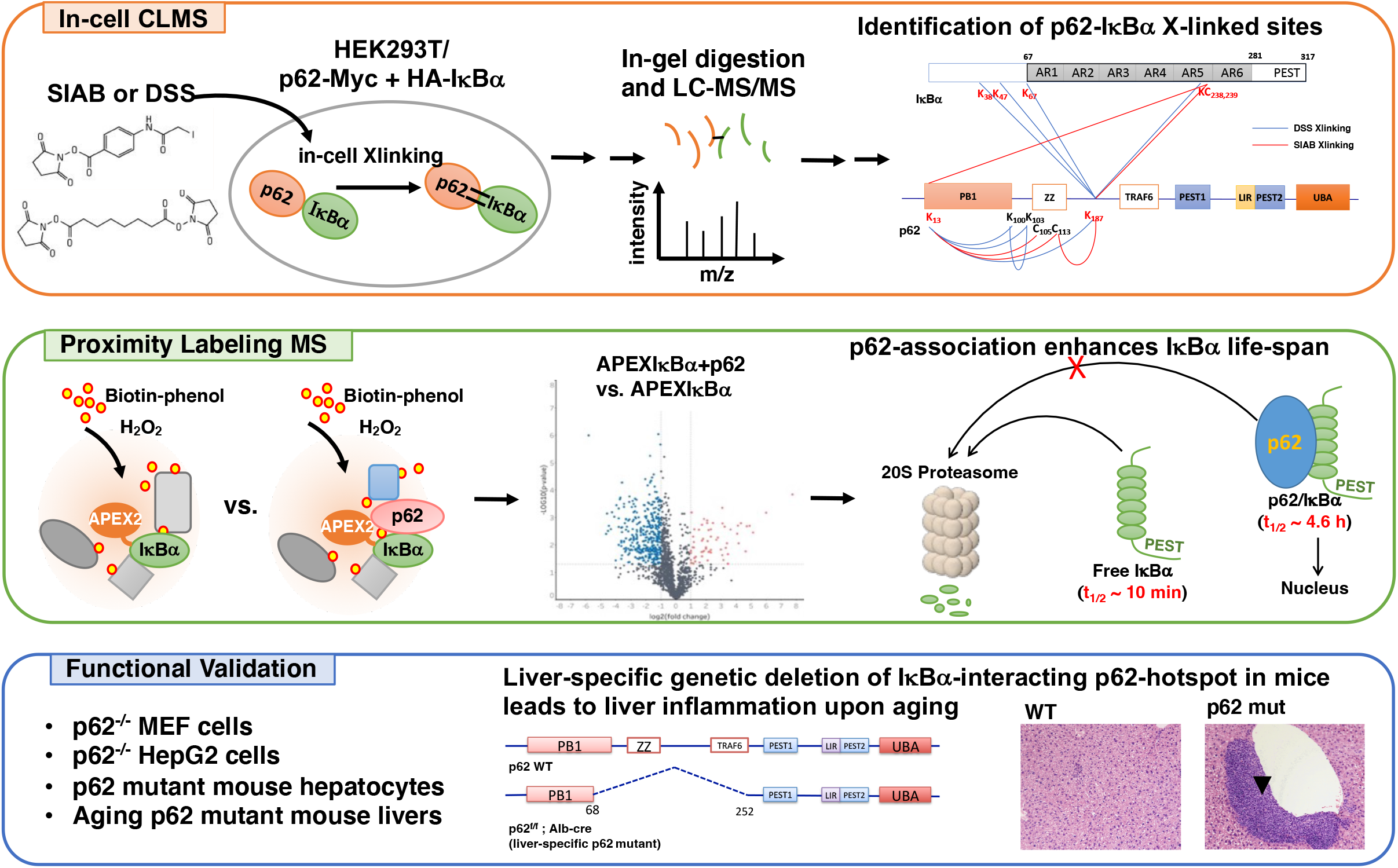

**Highlights:**

1. p62 binds to and stabilizes IκBα by preventing its proteolytic degradation
2. In-cell chemical crosslinking/LC-MS/MS identified the inter-crosslinked sites
3. Hotspots of p62-IκBα association are defined
4. APEX proximity labeling revealed p62 impaired IκBα-interaction with proteasome
5. p62 chaperones newly synthesized IκBα to terminate NF-κB activation.

**In Brief:** The transcriptional activator NF-κB inhibitor, IκBα is proteolytically unstable when uncomplexed. How newly synthesized IκBα escapes degradation to terminate nuclear NF-κB-activation is unknown. Using in-cell chemical crosslinking and proximity labeling MS analyses, we uncovered a novel association of p62 with IκBα via well-defined structural hotspots, which impairs its interaction with the 26S/20S proteasome, extending its life-span and enabling termination of NF-κB-activation. Mice carrying liver-specific genetic deletion of p62-IκBα hotspot exhibit enhanced liver inflammation upon aging, validating this novel p62 role.

## INTRODUCTION

The nuclear factor kappa-light-chain-enhancer of activated B cells (NF-κB) is a major transcriptional factor responsible for the nuclear activation of genes involved in myriad cellular processes, including inflammation, immune response and cancer. In the liver as in many cells, NF-κB exists largely as a p65/p50 heterodimer that is sequestered in the cytoplasm and kept transcriptionally inactive through its tight binding to specific NF-κB inhibitors (IκBs), which mask its nuclear localization signal (NLS) and DNA-binding domain (1). Of these IκBs, IκBα and IκBβ are the major isoforms in liver cells and cell-lines, each regulated by different extracellular signals and/or cues (2-5). Canonical transcriptional activation of NF-κB-responsive genes triggered by appropriate extracellular signals such as proinflammatory cytokines, entails the disruption of IκB-mediated NF-κB-tethering via IKK-elicited phosphorylation of IκB-N-terminal Ser_32/36_-residues that in turn triggers their ubiquitin (Ub)-dependent 26S proteasomal degradation (UPD) (6-8). Such IκB-destruction unleashes NF-κB, unmasking its NLS and enabling its nuclear translocation and subsequent DNA-association and transcriptional activation of NF-κB-responsive genes including those of IκBα (but not IκBβ) (9-12) and p62/SQSTM1 (13-16), as well as immune and inflammatory-response genes. This NF-κB-mediated transcriptional activation of the IκBα-gene results in *de novo* IκBα-synthesis which upon nuclear re-entry competes for the DNA-bound NF-κB, escorting it out of the nucleus into the cytoplasm, thereby aborting further NF-κB-activation and resetting the system to baseline (9, 11, 15, 16).

We have previously reported that in liver cells, the cellular multifunctional protein scaffold p62/SQSTM1 is closely associated with IκBα, co-aggregating with it upon Zn-protoporphyrin IX (ZnPP)-elicited protein aggregation (17). Their association, however, is not required for ZnPP-elicited IκBα-sequestration into insoluble aggregates resembling hepatic Mallory-Denk bodies (MDBs), as such ZnPP-elicited IκBα-aggregation is unscathed upon p62-knockout (KO) of mouse embryo fibroblasts (MEFs) as well as primary hepatocytes (17). Herein, we elucidate a novel feature of such an exquisite p62-IκBα intracellular interaction in the regulation of this IκBα/NF-κB-mediated vicious inflammatory cycle. We document that in contrast to that in corresponding wild types (WTs), p62^-/-^-MEF cells as well as CRISPR-Cas9-edited p62^-/-^ HepG2 cells exhibit not only a considerably blunted IκBα-feedback response (restoration) after TNFα- or IL-1β-stimulation, but also show a relatively more pronounced and prolonged nuclear p65-localization. More importantly, by using a novel in-cell chemical crosslinking mass spectrometry (CLMS) approach, we further identify a previously unrecognized p62-domain for IκBα-binding, whose direct interaction results in extending the physiological IκBα half-life (t_1/2_), and thus enabling its efficient termination of NF-κB activation. Accordingly, we document that liver-specific genetic deletion of this p62 IκBα-binding-domain in intact mice leads to severe hepatic inflammation upon aging, most likely by destabilizing IκBα and thereby disrupting the normal termination of NF-κB activation elicited upon *de novo* IκBα-synthesis. These findings attest to an exquisite role for p62 in the regulation of NF-κB-IκBα-mediated physiological and pathophysiological processes.

## EXPERIMENTAL PROCEDURES

### p62-Mutant (p62mut) Mice and p62-KO Cell lines

#### p62mut mice

We set out to generate p62 liver-conditional KO mice in C57BL/6N strain (C57BL/6N-Sqstm1^tm1a(KOMP)Wtsi^/Mmucd) with the assistance of UC Davis KOMP Facility starting from frozen embryos carrying Sqstm1/p62 tm1a KO allele. The KOMP Facility generated the tm1c heterozygous mice that were provided to us. We first bred these mice to obtain the homozygous p62 flp/flp tm1c mice that were in turn employed for Alb-Cre deletion to generate the liver-conditional p62 KO mice. For this purpose, Alb-Cre homozygous mice purchased from Jackson Laboratory (Bar Harbor, ME, strain 003574), were bred with the p62 flp/flp homozygous mice. The resulting pups, all p62 flp/+, Albcre/+ heterozygous, were then bred with the p62 flp/flp homozygous mice with 25% of the resulting pups being p62 flp/flp, Albcre/+, thus carrying specifically in the liver, a copy of Albcre recombinase to delete out the FLPed region in the p62 gene. After obtaining the p62 flp/flp, Albcre/+ conditional KO mice, p62 flp/flp, Albcre/+ mice were bred with p62 flp/flp mice, such that 50% of the pups would be p62 liver conditional KOs and the other 50% of pups, the WT controls. Immunoblotting (IB) analyses of their hepatic p62 content and RT-PCR followed by sequencing revealed however that instead of the anticipated liver-conditional p62 KO, a structural deletion mutant (p62mut), with the deletion of the region coding for amino acid residues 68-252, was obtained. Mice were fed a standard chow diet and maintained under 12-hour light/dark cycle. All animal experimental protocols were approved by the UCSF/Institutional Animal Care and Use Committee.

#### p62 KO MEF cells

Were generated from p62 KO mice by Prof. M. Komatsu’s lab (18), Niigata University, Japan, and provided by Prof. Haining Zhu, University of Kentucky.

#### p62 KO HepG2 cells

The CRISPR/Cas9 system was used to knockout p62 in HepG2 cell line. The CRISPR guide RNAs (gRNAs) were designed using CRISPR Design online tool (19). A sequence targeting exon 3 (TCAGGAGGCGCCCCGCAACA**TGG**) was used. Oligonucleotide containing the CRISPR target sequence was annealed and ligated into Bbs1 linearized pSpCas9(BB)-2A-Puro (PX459) vector (Addgene #48139). HepG2 cells were transfected with PX459-p62 with X-tremeGENE HP transfection reagent (Roche). Twenty-four h after transfection, cells were exposed to medium containing puromycin and selected for 48 h, after which, cells were recovered with an antibiotic-free medium for 24 h before re-seeding into 100 mm dishes at 100 cells/dish. Cells were allowed to grow for 4-8 weeks until single cell colonies appeared. Single cell colonies were then picked and expanded for screening using Western IB analyses as guidelines.

#### Cell culture and Transfections

p62mut mice and age-matched WT litter-mates (8-12-week old) bred in our lab were used for primary hepatocyte preparation. Hepatocytes were isolated by *in situ* collagenase perfusion and purified by Percoll-gradient centrifugation by the UCSF Liver Center Cell Biology Core, as described previously (20). Fresh primary mouse hepatocytes were cultured on Type I collagen-coated 60 mm Permanox plates (Thermo Scientific, Grand Island, NY) in William’s E Medium (Gibco, Grand Island, NY) supplemented with 2 mM L-glutamine, insulin-transferrin-selenium (Gibco, Grand Island, NY), 0.1% bovine albumin Fraction V (Gibco, Grand Island, NY), Penicillin-Streptomycin (Gibco, Grand Island, NY) and 0.1 μM dexamethasone. Cells were allowed to attach overnight and then overlaid with Matrigel by replacing with fresh media containing Matrigel (0.25 mg/mL; Corning, Oneonta, NY). From the 2nd day after plating, the medium was replaced daily, and cells were further cultured for 4-5 days with daily light microscopic examination for any signs of cell death and/or cytotoxicity. On day 5, some cells were treated with IL-1β (20 ng/mL) for 10 min, cells were then harvested at indicated timepoints using a cell scraper.

HepG2 cells were cultured in minimal Eagle’s medium (MEM) containing 10% v/v fetal bovine serum (FBS) and supplemented with nonessential amino acids and 1 mM sodium pyruvate. HEK293T and MEF cells were cultured in Dulbecco’s Modified Eagle high glucose medium (DMEM) containing 10% v/v FBS. For transfection experiments, cells were seeded on 6-well plates, when cells were 60% confluent, each cell well was transfected with 3 μg of plasmid DNA complexed with TurboFect transfection reagent (ThermoFisher, Grand Island, NY) for HEK293T cells, and X-tremeGENE HP transfection reagent (Roche, Indianapolis, IN) for HepG2 cells, according to the manufacturers’ instructions. At 40-72 h after transfection, cells were either treated as indicated or directly harvested for assays. For NF-κB activation assays, HepG2 cells or MEF cells were first rested in FBS-free medium for 4 h, then treated with TNFα (20 ng/mL) or IL-1β (20 ng/mL) for indicated times, after which cells were harvested using a cell scraper.

#### Plasmids

pSpCas9(BB)-2A-Puro (PX459) was a gift from Dr. Feng Zhang (Addgene plasmid # 48139; http://n2t.net/addgene:48139; RRID: Addgene_48139) (21). pCMV-3HA-IκBα and pCMV-3HA-IκBα-S_32_A/S_36_A were gifts from Dr. Warner Greene (plasmids #21985, #24143; Addgene, Cambridge, MA). pLX304-Flag-APEX2-NES were gifts from Dr. Alice Ting (Addgene plasmid # 92158; http://n2t.net/addgene:92158; RRID: Addgene_92158) (22). C1-Emerald was a gift from Dr. Michael Davidson (Addgene plasmid #54734). pcDNA6 and pcDNA3 vectors were from Invitrogen (Grand Island, NY). C1-Emerald-IκBα, pcDNA6-p62-myc were constructed in house previously (17). Various IκBα- and p62-mutants were constructed for this study. Truncation mutations were constructed by PCR cloning, or DNA fragment gene synthesis (Genewiz, NJ), or homologous recombination based NEBuilder® HiFi DNA Assembly kit (NEB, MA). Site-directed mutagenesis was carried out using QuikChange QuikChange Site-Directed Mutagenesis Kit (Agilent, CA). The primers, templates, vectors and restriction enzymes (RE) used are summarized below:

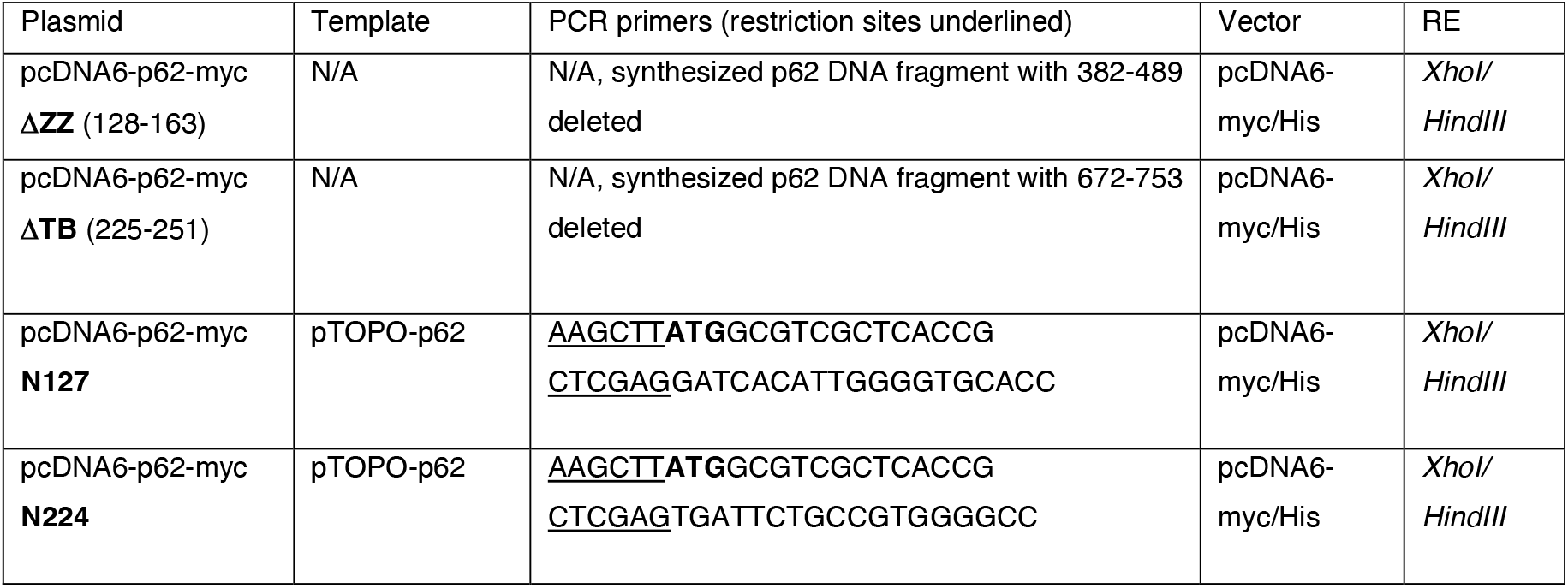
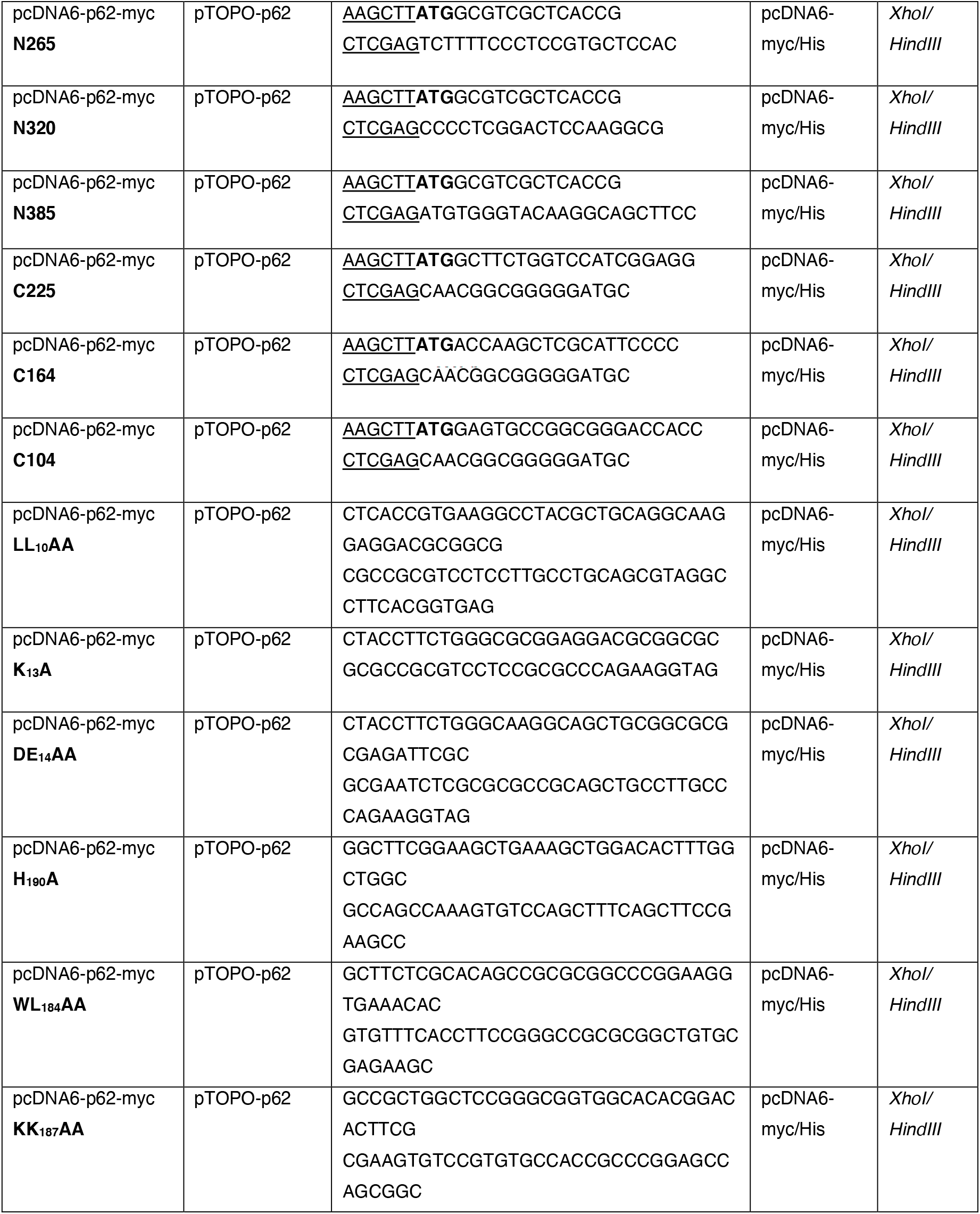
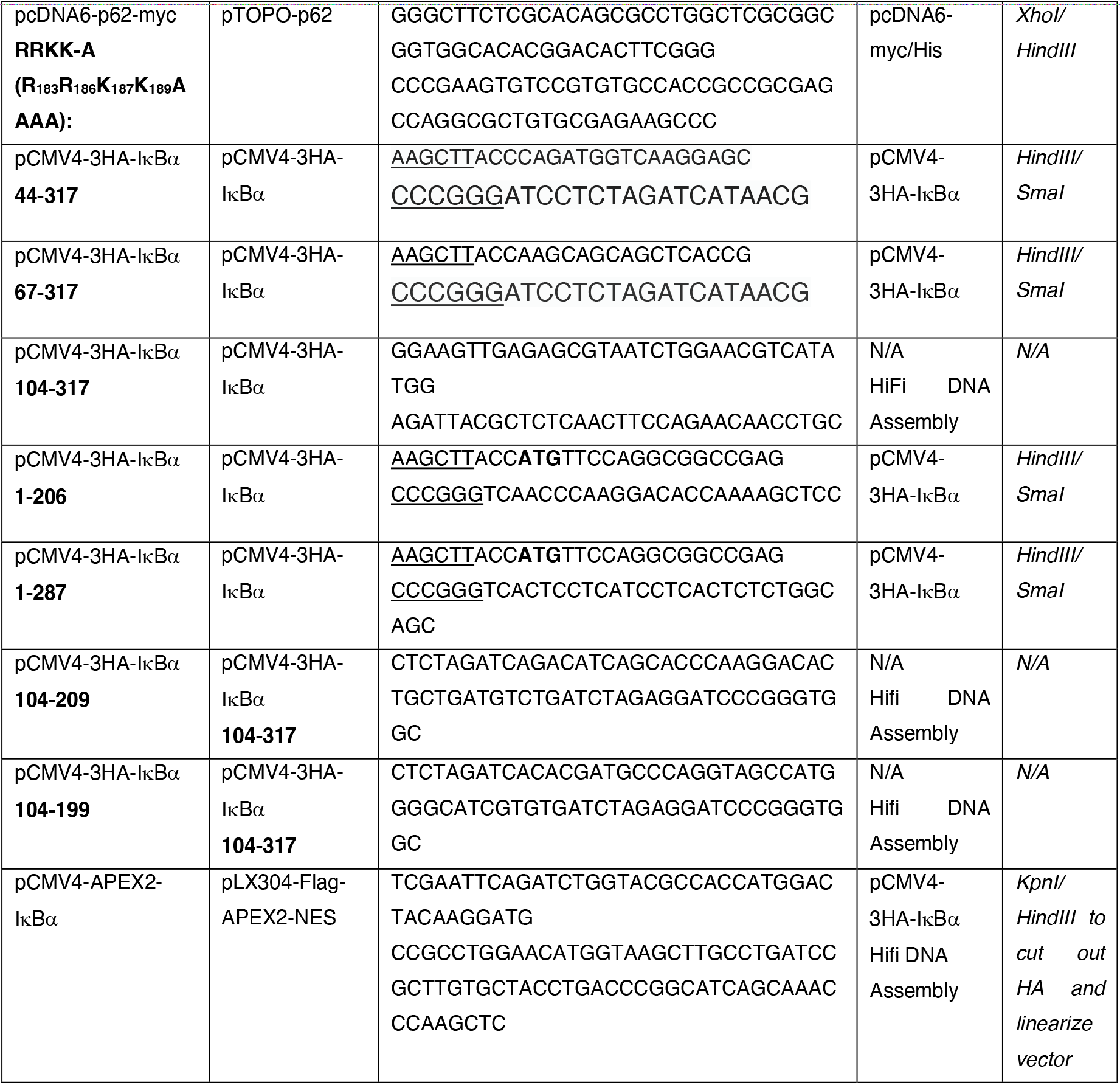

#### Cell Fractionation

For whole cell lysate preparation, cells were harvested in Cell Lysis Buffer (CSL buffer, Cell Signaling Technology, Danvers, MA) containing 20 mM Tris-HCl (pH 7.5), 150 mM NaCl, 1 mM EDTA, 1 mM EGTA, 1% Triton, 2.5 mM sodium pyrophosphate, 1 mM β-glycerophosphate, 1 mM Na_3_VO_4_, leupeptin (1 *μ*g/mL) and supplemented with 10% glycerol and a protease/phosphatase inhibitor cocktail (Pierce, Grand Island, NY). Cell lysates were sonicated for 10 s and then cleared by centrifugation at 4°C in a tabletop centrifuge at 14,000 g for 10 min. Nuclear and cytoplasmic extracts were prepared using NE-PER Nuclear and Cytoplasmic Extraction Kit (Thermo Scientific, MA). Briefly, cells were harvested in PBS supplemented with a protease/phosphatase inhibitor cocktail, and pelleted at 400 g for 5 min at 4°C. Cell pellet collected from one 60 mm petri dish was first resuspended in CER I buffer (400 μL) and incubated on ice for 10 min. Then CER II (22 μL) was added and the mixture vortexed, followed by incubation on ice for 1 min. The tubes were then centrifuged at 14,000g for 5 min at 4°C. The supernatants were then transferred into a new tube as cytoplasmic extracts. The tubes containing the pellet were then washed briefly with PBS and centrifuged briefly, and the residual supernatants were discarded. The pellets were then resuspended in ice-cold NER buffer (150 μL) and sonicated for 5 sec to disrupt the pellets. The tubes were then centrifuged at 14,000g for 5 min at 4°C and the supernatants were recovered as the nuclear extracts. Mitochondrial extracts were prepared using Mitochondria Isolation Kit for Cultured Cells (Thermo Scientific, MA) with some modifications. Briefly, cells from three 60 mm petri dishes were harvested in PBS supplemented with a protease/phosphatase inhibitor cocktail, and combined and pelleted at 400g for 5 min at 4°C. The pellets were then resuspended in Reagent A (600 μL) and incubated on ice for 2 min. Resuspended cells were then disrupted using a micro-tip homogenizer (Omni-Inc, GA) for 40 sec, twice. Reagent C (600 μL) was then added and mixed by inverting the tube several times. The tubes were then centrifuged at 700g for 10 min at 4°C. Nucleus and unbroken cells were pelleted at this step. The supernatants containing the cytoplasmic fraction were transferred to a new tube and then centrifuged at 12,000g for 20 min at 4°C. The supernatants (cytosolic fractions) were recovered and saved. The pellets were washed in Reagent C (200 μL) and then centrifuged at 12,000g for 5 min at 4°C. The supernatants were discarded and pellets were saved as the isolated mitochondria. Isolated mitochondria were then lysed in RIPA buffer (Cell Signaling Technology, Danvers, MA) supplemented with 0.1% SDS, 10% glycerol and a protease/phosphatase inhibitor cocktail with sonication. The pellet containing the nuclei and unbroken cells was first resuspended in Triton containing Cell Lysis Buffer with gentle pipetting to lyse unbroken cells but keeping the nuclei intact. The nuclei were then pelleted by centrifugation at 12,000g for 5 min at 4°C and the pellet was extracted using RIPA buffer with sonication, the resulting lysate was used as the nuclear extract.

#### Western Immunoblotting (IB) Analyses

For Western IB analyses, extracts from various cell fractions were prepared as described above. Protein concentrations were determined by the bicinchoninic acid (BCA) assay and equal amounts of proteins were separated on 4-15% Tris-Glycine eXtended (TGX) polyacrylamide gels. Proteins were transferred onto 0.2 μ nitrocellulose membranes (BioRad, Hercules, CA) for IB analyses.

The following primary antibodies were used: c-Myc (9B11, Cell Signaling Technology, Danvers, MA), HA (C29F4, Cell Signaling Technology, Danvers, MA), IκBα (C-terminus, 44D4, Cell Signaling Technology, Danvers, MA), Phospho-IκBα (Ser_32_) (14D4, Cell Signaling Technology, Danvers, MA), p62 (2C11, Abnova, Taipei City, Taiwan), NF-κB p65 (D14E12, Cell Signaling Technology, Danvers, MA), Phospho-NF-κB p65 (93H1, Cell Signaling Technology, Danvers, MA), β-Actin (Sigma, St. Louis, MO), Histone H3 (Abcam, Cambridge, MA), COX IV (3E11, Cell Signaling Technology, Danvers, MA).

#### Co-Immunoprecipitation (Co-IP) analyses

Whole-cell extracts were prepared as described above. Cell lysates (1 mg) were then incubated with EZview™ Red Anti-HA Affinity Gel (30 μL; MilliporeSigma, MA) or ChromoTek Myc-Trap® Agarose (30 μL; Proteintech, IL) at 4°C overnight, and then eluted by heating at 70°C for 10 min in 2X SDS-loading buffer. Eluates were subjected to IB analyses as described above.

#### In-Cell Chemical Crosslinking and crosslinked protein capture for LC-MS/MS (CLMS)

For each crosslinking, ten 100 mm petri dishes of cultured HEK293T cells were co-transfected with pCMV4-3HA-IκBα and pcDNA6-p62-myc at a 1: 1 ratio for 48 h. For DSS crosslinking, cells were treated with 10 μM NEM for 5 min right before harvesting, Cells were then washed 3 times using ice-cold PBS and collected by a cell scraper in PBS. After that, DSS (disuccinimidyl suberate; 0.05 mM) or SIAB [succinimidyl (4-iodoacetyl)aminobenzoate); 0.05 mM] was added directly to the cell suspension and mixed gently by inversion and incubated on a rocker platform for 20 min in the dark. Tris buffer pH 7.4 was then added to a final concentration of 20 mM to quench the crosslinking reaction. Cells were pelleted and then lysed in RIPA buffer supplemented with 10 μM NEM with sonication and cleared by centrifugation. The supernatant was then incubated with Myc-Trap (500 μL) overnight at 4°C to pull-down Myc-tagged p62 and HA-IκBα-crosslinked Myc-tagged p62. Beads were then washed three times using RIPA buffer and then eluted by heating at 70°C for 10 min in 2X SDS-loading buffer containing 4% SDS. Eluates were then diluted 20 times using CSL buffer and incubated with HA-agarose (100 μL) overnight at 4°C. Subsequently, beads were collected by centrifugation at 3,000g for 30s and then washed 3 times with RIPA buffer and then eluted by heating at 70°C for 10 min in 2X SDS-loading buffer. This tandem IP resulted in an enrichment of p62-Myc and HA-IκBα crosslinked species. Eluates were then split into 4-5 gel lanes and subjected to SDS-PAGE and stained with Coomassie Blue to visualize the bands for subsequent in-gel digestion.

#### IR fluorescence detection of DSS- or SIAB-crosslinked p62-IκBα species

Western IB analysis of the crosslinked p62-IκBα complexes was first carried out with a two-color system using IRDye® 680RD goat anti-mouse IgG for labeling p62-Myc and IRDye® 800CW goat anti-rabbit IgG for HA-IκBα, which were then detected by IR fluorescence detection with an Odyssey® Fc Imaging System (LI-COR Biosciences).

#### APEX reaction and biotinylated protein capture

APEX biotinylation was carried out as described by Hung, et al. (22). Briefly, HEK293T cells grown in 60 mm petri dishes were transfected with N-terminal pCMV4-APEX2-IκBα with or without pcDNA6-p62-Myc co-transfection. mEmerald (GFP)-tagged IκBα (GFP-IκBα) co-transfected with pcDNA6-p62-Myc were used as a negative control. Forty-eight h after transfection, cells were treated with 500 μM biotin-phenol for 30 min. H_2_O_2_ was then added to a final concentration of 1 mM for 1 min to initiate the catalytic biotinylation labeling. The labeling reaction was then stopped by quickly exchanging the media with PBS containing 500 mM sodium ascorbate and 5 mM sodium azide. Cells were further washed 2 times with the antioxidant containing PBS buffer and then lysed in RIPA buffer. Cell lysates were cleared by centrifugation at 14,000g, the supernatants were screened by Western IB analyses with streptavidin-HRP to determine the extent of the biotinylation, with subsequent streptavidin (SA) pull-downs to enrich the biotinylated proteins. For SA pull-downs, Pierce™ Streptavidin Magnetic Beads (75 μL; ThermoFisher, MA) were added to the lysates and the mixture incubated at 4°C overnight on a rotator. Beads were then washed twice with RIPA buffer, once with 1 M KCl, once with 2 M urea in 20 mM Tris base, twice with RIPA buffer and lastly twice with 100 mM ammonium bicarbonate (ABC) before proceeding with the on-bead digestion.

#### Mass spectrometry (MS)

For in-gel digestion of *in cell* crosslinked samples, the indicated gel bands from 4 or 5 replicate lanes were pooled and processed using a standard in-gel digestion procedure (23, 24). Briefly, each gel piece was excised into 1mm x 1mm pieces, reduced and alkylated, and then digested overnight with 300 ng of Trypsin/Lys-C Mix (Promega, Madison, WI). For on-bead digestion following SA pull-down, beads were first resuspended in 6 M urea, 100 mM ammonium bicarbonate (ABC) buffer, and then reduced by addition of 10 mM (final) DTT and incubation at 37°C for 30 min. The samples were then alkylated with 15 mM (final) iodoacetamide at room temperature in the dark for 30 min. Trypsin/Lys-C Mix (100 ng) was then added, and the mixture incubated for 4 h at 37°C. Samples were diluted 1:6 (v:v) with 100 mM ABC to reduce the urea concentration to 1 M and then further incubated at 37°C overnight. The resulting peptide mixture was desalted using C18-Zip tips (EMD Millipore, Billerica, MA), speed-vacuumed to dryness and suspended in 0.1% formic acid for injection into: a Q-Exactive Plus (for APEX experiments), an LTQ-Orbitrap Velos (for SIAB cross-linking), or an Orbitrap Fusion Lumos (for DSS cross-linking) mass spectrometer (Thermo Fisher, San Jose, CA) coupled through an EASY-Spray nano ion source (Thermo) to a nano-Acquity UPLC (Waters, Milford, MA) running an EASY-Spray column (75 μm x 15 cm column packed with 3 μm, 100 Å PepMap C18 resin; Thermo). The mobile phases were: solvent A water/0.1% formic acid; solvent B: acetonitrile/0.1% formic acid. For Apex analyses, the samples were loaded at 600 nL/min of 2% B, followed by elution at 400 nL/min with a gradient to 23% B over 103 min and then a second gradient to 40% B in 11 min followed by washing at 75% B and re-equilibration. Total run time was 150 min. Precursor ions were acquired from 350-1500 m/z (70k resolving power, 3e6 AGC, 100 ms max inj. time), the top 10 doubly charged or higher precursor ions were selected with a 4 m/z window, dissociated with 25% NCE HCD, and measured at 17.5k resolving power (5e4 AGC, 120 ms max inj. time). A 15 sec dynamic exclusion window was used.

DSS-crosslinked samples were loaded at 600 nL/min at 2%B, and then eluted with a 400 nL/min gradient from 3-27% B over 92 min. The column was then washed at 75% B and re-equilibrated back to 2%. The total run time was 120 min. Precursor ions were acquired from 375-1500 m/z in the Orbitrap (120k resolving power, 4e5 AGC, 50 ms max inj. time). The top 9 precursors with charges 3-8+ and intensity greater than 50,000 were isolated in the quadrupole (1.6 m/z selection window) and sequentially dissociated by HCD with stepped 25, 30, 35% NCE and EThcD (25 ms ETD reaction time with 10% supplemental HCD activation). Product ions were acquired in the Orbitrap (30k resolving power, 1e5 AGC, 150 ms max inj time). 30s dynamic exclusion and the peptide monoisotopic ion precursor selection option were enabled.

SIAB-crosslinked samples were loaded at 600 nL/min 3% B, and then eluted at 300 nL/min with a gradient from 3-28% B over 90 min. The column was then washed at 75% B and re-equilibrated with a total run time of 125 min. Precursor ions were acquired in the Orbitrap from 300-1800 m/z (30k resolving power, 2e6 AGC, 100 ms max inj. time). The top 6, 3+ and higher precursors with intensity greater than 4000 were isolated in the quadrupole (4 m/z isolation window) and dissociated by HCD (30% NCE). Product ions were measured in the Orbitrap (7.5k resolving power, 9e4 AGC, max inj time 500 ms).

For Apex analyses, peaklists were extracted using PAVA, an in-house software developed by the UCSF Mass Spectrometry Facility. Peaklists were searched against the SwissProt human database (SwissProt.2019.4.8; 20418 entries searched) using ProteinProspector (version 5.19.1; http://prospector.ucsf.edu/prospector/mshome.htm) (25). A fully randomized decoy database (an additional 20418 entries) was used to estimate false discovery rates (FDR) (26). Protein Prospector search parameters were as follows: tolerance for precursor and product ions were 20 ppm and 30 ppm, respectively; a maximum of 1 missed cleavage of trypsin was allowed; carbamidomethylation of Cys was set as a fixed modification; variable modifications were set to: N-terminal Met loss and/or acetylation, Met oxidation, peptide N-terminal Gln to pyroGly conversion; the maximum number of variable modifications was 2. Reporting thresholds were as follows: Minimum score of protein: 22.0; minimum score of peptide: 15.0; maximum E value of protein: 0.01; maximum E value of peptide: 0.05. These score thresholds resulted in a <1% FDR at the peptide level and 5% FDR at the protein level.

For crosslinking analyses of the double affinity purified IκBα, p62 pulldowns, peaklists corresponding to the excised gel bands were initially searched to identify the most abundant proteins in the sample. As expected, p62 and IκBα were detected at an order of magnitude greater abundance measured by normalized spectral abundance factor (NSAF) (27) than other endogenous (ie. not obviously artifactual) human proteins (**Table S4**). A restricted database consisting of all proteins detected with more than one unique peptide and NSAF values within two orders of magnitude of the most abundant protein was used in the crosslinking search (DSS: 28 sequences total, SIAB: 25 sequences. **Table S4** protein IDs: purple accession numbers). Randomized and 10x longer versions of the target sequences were used for FDR control and scoring model training. Peaklists were searched with Protein Prospector v6.4.23 with Trypsin specificity and 2 missed cleavages. Precursor and product ion tolerance were 6 and 12 ppm respectively. DSS- or SIAB-crosslinking was specified with 5,000 intermediate hits saved and the rank of the best peptide was restricted to 1. For DSS experiments, NEM modification of Cys was used as a constant modification. For SIAB experiments, no constant modification was specified and carbamidomethylation of Cys was included as a variable modification. For both experiments, variable modifications were: Met oxidation, loss and/or acetylation of protein N-terminal Met, peptide N-terminal Glu conversion to pyroglutamate, dead-end DSS modification at Lys and protein N-terminus or dead-end SIAB modification of Cys, and incorrect monoisotopic peak assignment (neutral loss of 1Da). Up to 3 variable modifications per peptide were allowed. Crosslinked matches were classified using Touchstone (in-house R package, *manuscript in development*) with a minimum peptide length of 3 amino acids and minimum score difference of 0. Crosslinks were reported at the unique-residue pair level with 1% FDR. When crosslinks could not be uniquely identified all possible hits were reported (**Table S4**).

#### Experimental Design and Statistical Rationale

The goal of the IκBα-APEX proteomic experiments was to identify proteins that interact with IκBα and the changes in these interactions upon p62 co-expression. We first used the SAINTexpress software package (28) to independently identify proteins specifically labeled by APEX2-IκBα with or without p62-Myc co-expression using GFP-IκBα (C1-IκBα) with p62-Myc co-expression as the negative control. Four biological replicates were performed for each condition. The Protein Prospector search results (**Table S1**) were reformatted and analyzed by SAINTexpress on the CRAPome website (www.crapome.org) (29). The SAINT output (**Table S2**) was first filtered by BFDR (Bayesian False Discovery Rate) score, proteins with a BFDR score below a threshold of 0.01 in either APEX2-IκBα with p62-Myc or APEX2-IκBα without p62-Myc co-expression were retained. Average fold change values were calculated from the spectral counts of APEX2-IκBα + p62-Myc vs. APEX2-IκBα alone and p-values were calculated by a t-test, to generate the volcano plot (**Fig. 6B**). Selected proteins that showed a significant change upon p62-Myc co-expression were searched using MaxQuant (30) for more accurate label-free intensity-based quantification (**Table S3**). The intensities were then normalized using IκBα intensities in each sample to generate the bar charts (**Fig. 6C-G**).

## RESULTS

### Intimate protein-protein interactions of p62 with IκBα enhances its proteolytic stability

Co-transfection of HEK293T cells with Myc-tagged p62 (p62-Myc) plasmid coupled with HA-tagged IκBα (HA-IκBα) plasmid, and subsequent anti-Myc IgG-immunoprecipitation of the cell lysates led to the co-immunoprecipitation of HA-IκBα, attesting to an intimate intracellular interaction of the two proteins (**Fig. 1A**). We also found that co-transfection of the two plasmids at 1 μg each resulted in a much greater expression of IκBα-protein than that observed after 2 μg-transfection of HA-IκBα plasmid alone. These findings suggested that upon co-expression, p62 could be either enhancing IκBα-synthesis or protecting IκBα from degradation. To directly assess the role of p62 in IκBα-protein stability, we carried out cycloheximide (CHX) pulse-chase analyses in HEK293T cells either expressing HA-IκBα alone or co-expressing both HA-IκBα and p62-Myc. p62-coexpression indeed extended the cellular IκBα t_1/2_ from 0.25 h under basal conditions to a t_1/2_ of 4.57 h, i.e. an 18-fold life-span prolongation (**Fig. 1B**). This revealed that p62 enhanced IκBα cellular stability by protecting it from proteolytic turnover. Intracellular IκBα-turnover involves various cellular proteolytic systems, among them the 20S/26S proteasomal and calpain systems being the most prolific under basal conditions (31-33). Thus, upon expression of IκBα alone, it was largely protected by the proteasomal inhibitor MG-132, and to a lesser extent by the calpain-inhibitor, calpeptin (**Fig. 1C**). By contrast, p62-co-expression with HA-IκBα had a major protective role that tended to minimize the additional contributions of these protease inhibitors to IκBα-stability (**Fig. 1C**). Moreover, the near comparable IκBα-levels upon p62-coexpression in the presence or absence of MG132 suggested that the IκBα-enhancement was not due to its p62-mediated enhanced synthesis. These findings conclusively indicated that p62-interaction affords cellular protection against rapid IκBα proteolytic turnover.

**FIGURE 1.**
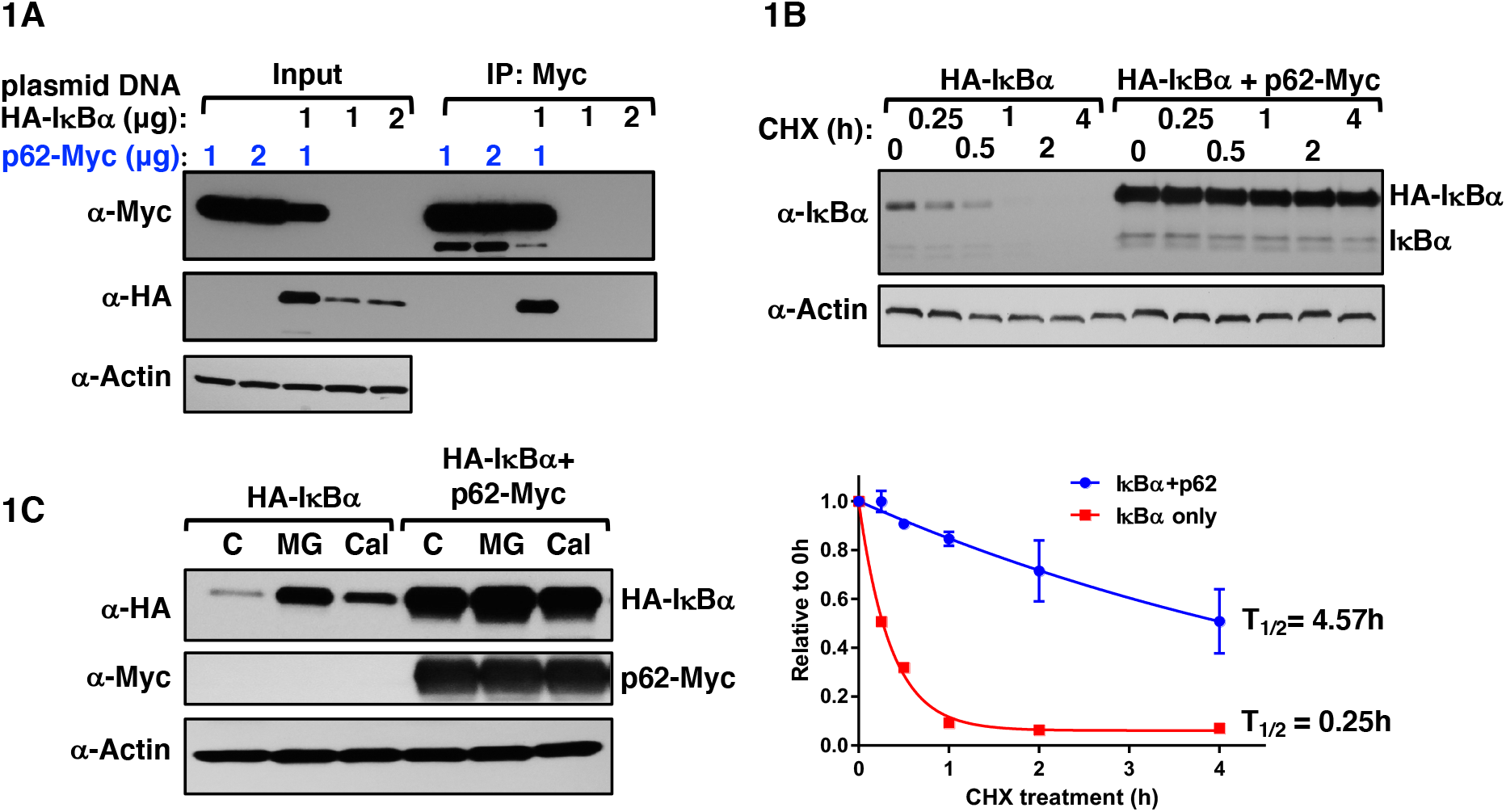
p62 stabilizes IκBα protein by blocking its proteolytic turnover. **A**. HEK293T cells were seeded in 6-well plates overnight, and then each well was transfected with indicated amounts of pCMV4-3HA***-***IκBα vector (1 or 2 μg) or co-transfected with pcDNA6-p62-Myc (1 or 2 μg). Whenever necessary, pcDNA6-Myc empty vector was used to supplement such that the total plasmid DNA amount transfected was 2 μg. Forty-eight h after transfection, cell lysates (10 μg) were used for IB analyses with actin as the loading control. **B**. HEK293T cells were transfected either alone with pCMV4-3HA***-***IκBα or co-transfected with pcDNA6-p62-Myc (1μg) as in (**A**). Forty-eight h after transfection, cells were treated with cycloheximide (CHX; 50 μg/mL) for indicated times, and cell lysates (10 μg) subjected to IB analyses with actin as the loading control. IκBα amounts relative to 0 h control were quantified from 3 experimental replicates and plotted. The half-life (t_1/2_) of IκBα with or without p62 co-expression was calculated based on a single exponential fit of the data with Prism Graphpad Version 6.07. **C**. HEK293T cells were transfected either alone with pCMV4-3HA***-***IκBα (1μg) or co-transfected with pcDNA6-p62-Myc (1μg) as in (**B**). Forty-two h after transfection, cells were treated with vehicle control (C), or MG132 (MG; 20 μM) or calpeptin (Cal; 100 μM) for 6 h, cell lysates (10 μg) were used for IB analyses with actin as the loading control.

### Identification of the p62-IκBα-interaction domains through structural deletion, in-cell chemical crosslinking/LC-MS/MS and site-directed mutagenesis analyses

To identify the p62-subdomains involved in its IκBα-interaction, we carried out p62-deletion analyses through sequential deletion of various p62-plasmid regions corresponding to structural elements that interact with various cellular protein partners and/or motifs [Phox and Bem1p-domain (PB1), Zn-finger binding motifs (ZZ), TRAF6-binding (TB), LC3-interacting region (LIR), PEST1 and PEST2 motifs, Keap1-interacting region (KIR), and Ub-association region (UBA)] (15 *and references therein;* 34-40) (**Fig. 2**). Each of these Myc-tagged plasmids including the WT p62-Myc plasmid was co-transfected along with the HA-IκBα-plasmid into HEK293T cells, followed by p62-Myc-immunoprecipitation and Western immunoblotting analyses of the co-immunoprecipitated HA-IκBα. These sequential deletion analyses identified the minimal p62-subdomain required for cellular IκBα-interaction and stabilization as the N-terminal 1-224 (N1-224) p62 residues that contained the PB1, ZZ and the intervening region (IR) between the ZZ- and TB-subdomains (**Fig. 2**). The C-terminal domain beyond the TB-subdomain was not required for this interaction.

**FIGURE 2.**
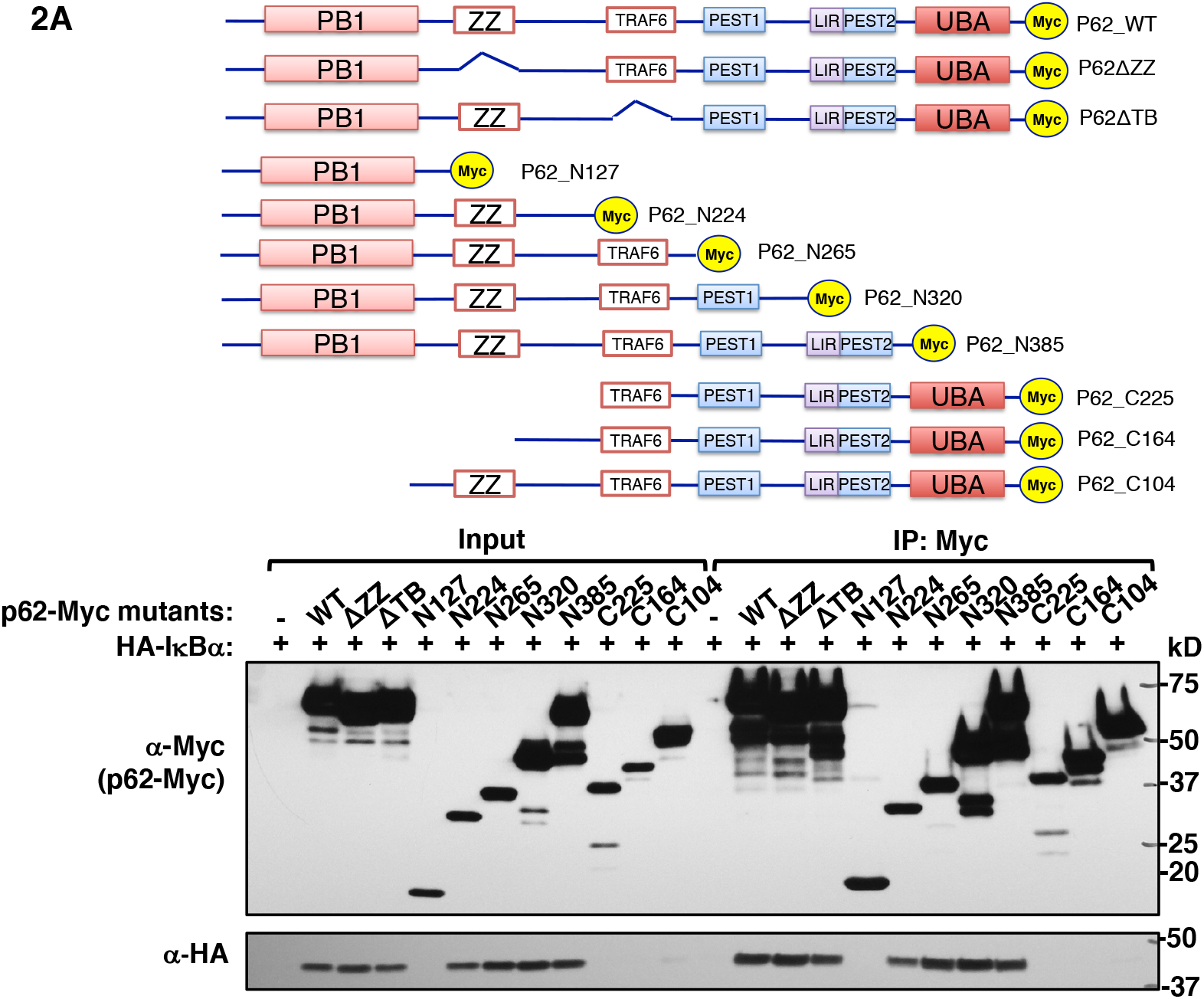
p62-PB1 domain and -linker between ZZ- and TRAF6-domains are both important for IκBα stabilization. p62 deletion and truncation mutations were constructed as schematically shown. HEK293T cells were seeded in 6-well plates overnight, and then each well was co-transfected with pCMV4-3HA***-***IκBα (1μg) and either WT or indicated mutant pcDNA6-p62-Myc vector (1μg) for 48 h. Whole cell lysates (500 μg) were used for co-IP with Myc-trap (50 μL), followed by IB analyses with HA- and Myc-antibodies.

To identify the hotspots of p62-mediated IκBα-recognition within this N1-224 subdomain under physiological conditions, we carried out in-cell chemical-crosslinking of HEK293T cells overexpressing HA-IκBα and p62-Myc using a cell-permeable chemical crosslinker either to crosslink amines to sulfhydryls, SIAB (0.05 mM), or amines to amines, DSS (0.05 mM) (**Fig. 3A**). The cells were lysed with denaturing RIPA buffer to disrupt native (non-crosslinked) protein-protein interactions. To obtain highly enriched crosslinked p62-IκBα species, the cleared RIPA lysates of DSS- or SIAB-treated cells were subjected to a dual immunoprecipitation: First with Myc-trap to pull-down p62-Myc, and subsequently with anti-HA-agarose. The tandem immunoprecipitated eluates along with an aliquot of their parent lysate were then subjected to SDS-PAGE (**Fig. 3C**), two-color Western immunoblotting of p62 and IκBα species followed by IR fluorescence detection (**Fig. 3B**). Overlap of the immunoblot fluorescence signals detected with either IRDye® 680RD goat anti-mouse IgG (p62-Myc) or IRDye® 800CW goat anti-rabbit IgG (HA-IκBα) enabled the visualization of the SIAB-crosslinked p62-IκBα species as yellow bands with molecular masses corresponding to the heterodimeric (1:1) = 150 kDa species, as well as additional heterooligomeric crosslinked species at higher (>150 kDa) molecular masses (**Fig. 3B**). Similar analyses were also carried out with DSS (**Fig. 3D; Fig. S1**). The gel-bands corresponding to these crosslinked p62-IκBα species were excised and subjected to in-gel digestion and LC-MS/MS analyses of the resulting crosslinked peptides. While p62 and IκBα were by far the most abundant endogenous proteins detected in these gel bands, potential crosslinks were searched against all proteins detected in the top two orders of magnitude with respect to abundance. A number of p62 crosslinks were discovered on a short peptide ^187^KVK^189^. Due to its length, it is difficult to unambiguously assign these crosslinks to p62 rather than to alternative sequences. For instance, hnRNP R peptide ^370^VKK^372^ in the DSS bands can only be distinguished by a single y_2_-ion and UBA52 peptide ^126^KVK^128^ in the SIAB bands is identical to the p62 sequence. All redundant explanations of these crosslinks are listed in **Table S4** and the corresponding spectral annotations are available online. However, we note that the crosslinked peptides originate from gel bands which have been doubly selected for p62-IκBα interactions and further, that p62 is detected at ∼20 and ∼100x higher NSAF levels than UBA52 and hnRNP R in these bands. Furthermore, only peptides from the Ub-domain of UBA52 were otherwise detected, while the KVK peptide is at the C-terminus of UBA52. Therefore, it is likely that p62 protein inference is correct for crosslinked peptides containing this sequence, as subsequently verified through our site-directed mutagenesis analyses of this p62-subdomain (*see below*). Two SIAB-crosslinked peptides were detected (**Table 1**), one between IκBα-Cys_239_ in its 5^th^ ankyrin-repeat (AR5) and p62-K_13_ in its PB1-subdomain, and the other between IκBα-Cys_239_ and p62-K_187_ in the intervening region (IR) between its ZZ- and TB-subdomains, consistent with the SIAB-ability to crosslink Lys- and Cys-residues (**Fig. 3D; Fig. S2A**). While similar DSS-crosslinked p62-IκBα species were also found, we found that the detection of these DSS-crosslinked species was greatly enhanced upon intracellular pretreatment with NEM (10 μM; 5 min) before cell-harvest, followed by NEM (10 μM) addition to the lysates (**Fig. S3D, E)**. With this NEM pretreatment, the optimal DSS in-cell crosslinking concentration was found to be 0.05 mM, just 1% of that recommended in the manufacturer’s instructions. (**Fig. S3E**). Both NEM-pretreatments greatly enhanced p62-IκBα-interactions (**Fig. S3A)**. Most likely, such NEM-elicited enhancement of DSS-mediated crosslinking was plausibly due to its maximizing available free K-residues by preventing their intracellular ubiquitination and/or sumoylation. Indeed, intracellular NEM-treatment consistent with its SH-reactivity, completely aborted all cellular protein ubiquitination and much of the nuclear sumoylation, further enhancing these p62-IκBα interactions (**Fig. S3B**). NEM was however incompatible with SIAB-crosslinking due to its well-recognized Cys/SH-reactivity (**Fig. S3D**).

**Table 1.**
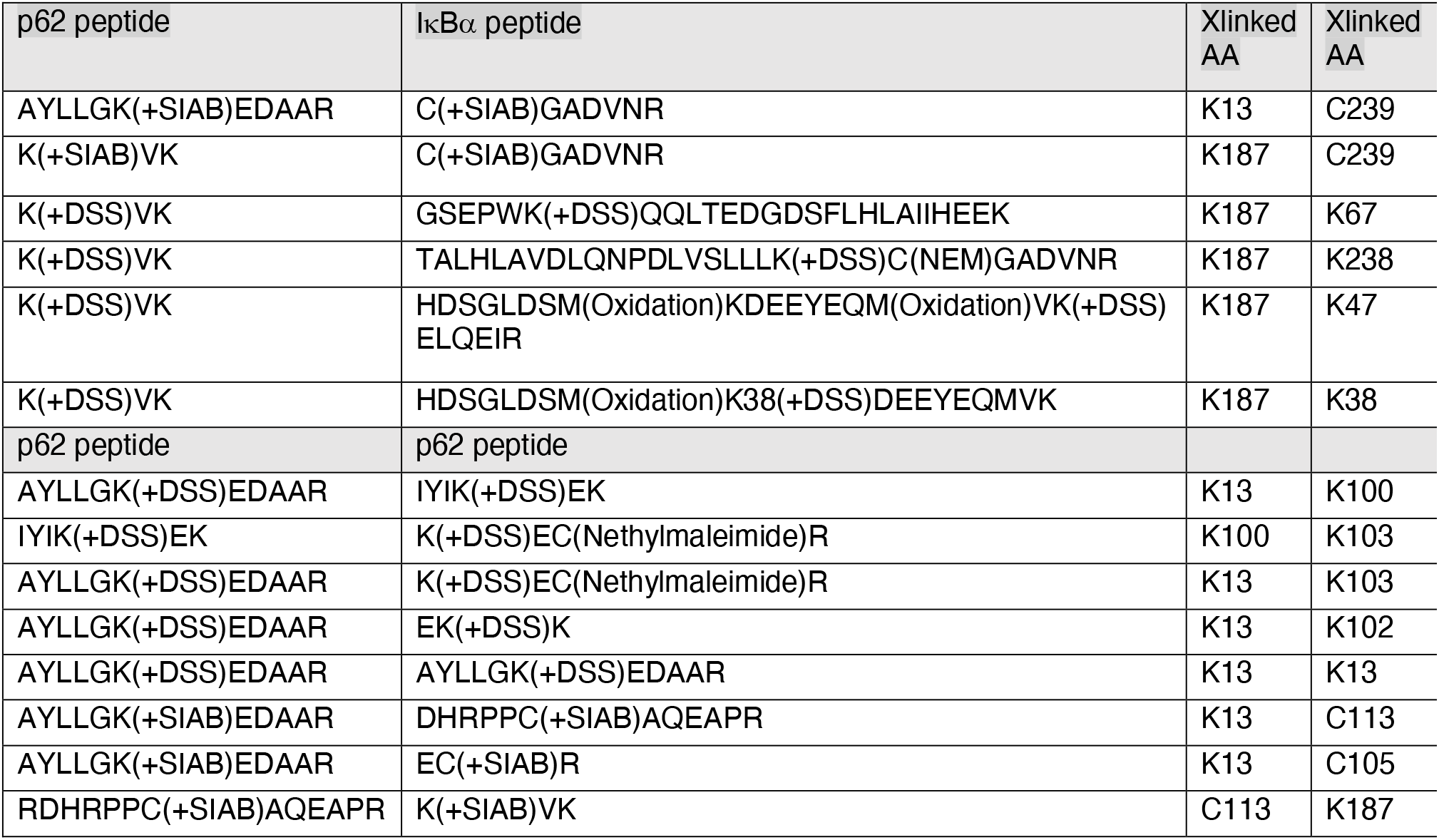
Crosslinked p62-IκBα peptides detected upon LC-MS/MS analyses.

**FIGURE 3.**
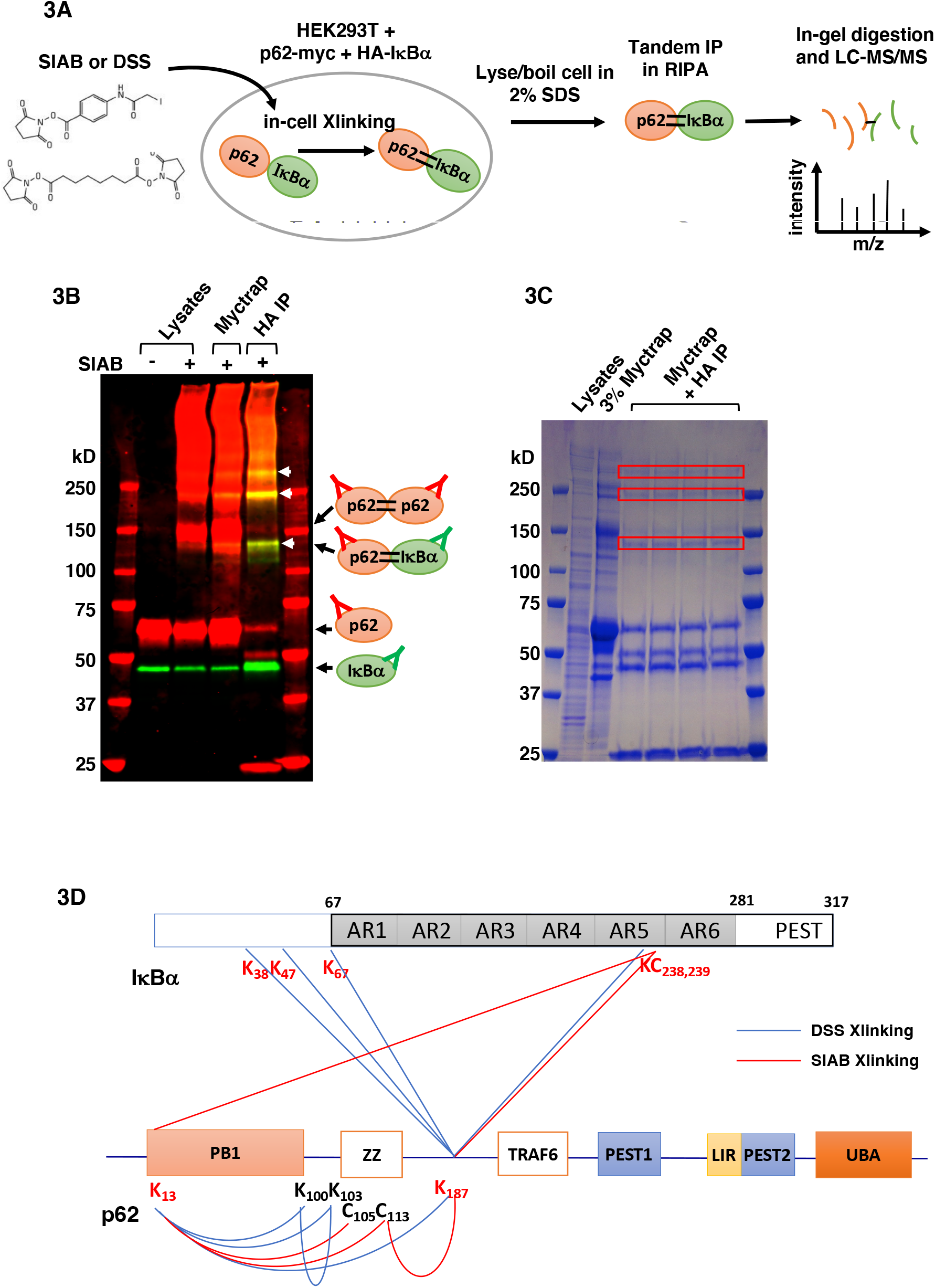
Potential p62-IκBα interaction hotspots identified by in-cell chemical crosslinking and mass spectrometric analyses. **A**. Workflow for p62-IκBα in-cell chemical crosslinking for mass spectrometric analyses. **B**. Representative Western IB analyses of p62-Myc and HA-IκBα after in-cell chemical crosslinking and tandem IP, first with Myc-trap followed by HA-agarose. Crosslinked species were enriched after tandem IP, as indicated by white arrows. **C**. Representative SDS-PAGE from large scale in-cell chemical crosslinking and tandem IP. Bands excised for in-gel digestion and LC-MS/MS are indicated by the red boxes. **D**. Summary of inter- and intra-crosslinked sites between p62-IκBα upon DSS- and SIAB-mediated crosslinking.

Similar in-gel digestion and LC-MS/MS analyses of the enriched immunoprecipitated DSS-crosslinked p62-IκBα species yielded four DSS-crosslinked peptides (**Table 1**): All these peptides documented interactions of K_187_ in the p62 IR with various IκBα K-residues residing in its N-terminus (K_38_, K_47_, K_67_) as well as K_238_, contiguous to the previously identified SIAB-crosslinked C_239_ in its AR5 (**Fig. 3D**). Co-expression of an IκBα mutant K_238_R with p62-Myc stabilized the basal IκBα-p62 interaction over that with the WT IκBα, and this interaction was further enhanced when an IκBα mutant (9KR) with all its 9 K-residues mutated to R, was co-expressed (**Fig. S3C**). These findings suggest that (i) a positively charged IκBα K- or R-residue is involved in these interactions, and (ii) in these p62-IκBα interactions, an R-residue is apparently favored over K, plausibly by circumventing post-translational K-modifications that would disrupt these interactions.

Together, the above CLMS analyses reveal that the N-terminal p62 domain (N1-224) forms important interactions with two IκBα-subdomains: One in its N-terminus (encompassing K_38_, K_47_, and K_67_) and the other in its C-terminal AR5 (around K_238_/C_239_). The latter site is in close proximity to AR6 (residues 243-280), which apparently harbors a degron (residues 251-262) for Ub-independent proteasomal degradation of IκBα (32).

To further define the importance of these p62-subdomains to its IκBα-interaction, we carried out site-directed mutagenesis of p62-residues flanking K_13_ and K_187_ followed by a HEK293T-cell co-transfection of Myc-tagged constructs and an anti-Myc-immunoprecipitation approach similar to that used above (**Fig. 2**). These findings revealed that mutation of p62-residues L_10_/L_11_, K_13_ or D_14_/E_15_ to Ala had very little influence on its IκBα-protein interactions (**Fig. 4**). On the other hand, while mutation of p62-residues W_184_/L_185_ and K_187_/K_189_ to ALA reduced its IκBα-protein interactions, that of H_190_ had little effect. By contrast, mutation of p62-residues R_183_/R_186_/K_187_/K_189_ flanking its IκBα-crosslinked K_187_-site completely aborted this interaction (**Fig. 4**), thereby identifying this positively charged p62-microregion as a hotspot for IκBα-interaction.

**FIGURE 4.**
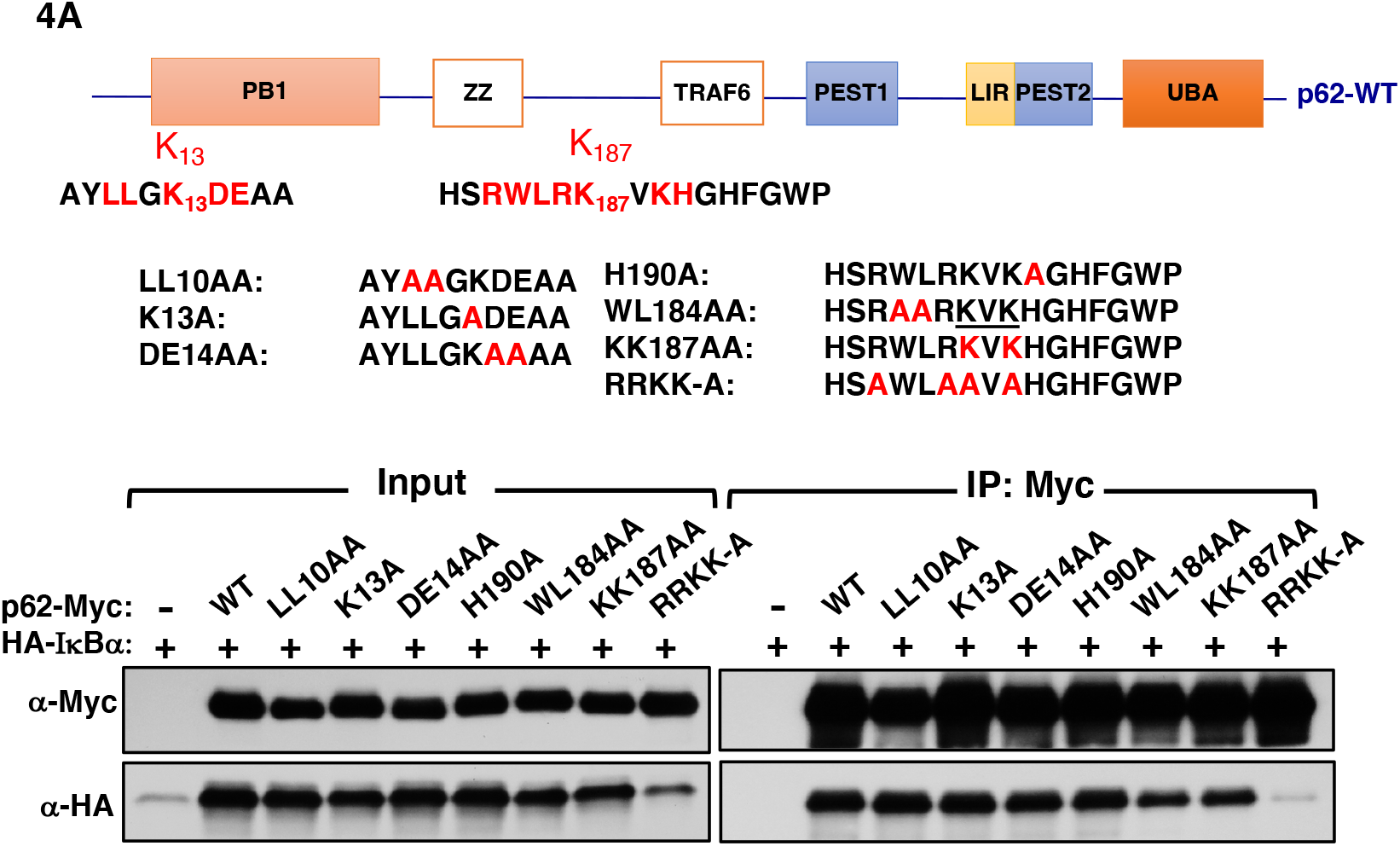
p62-IκBα interaction sites identified by site-directed mutagenesis. Potential sites in p62 identified as IκBα-interaction hotspots (**Fig. 3**) were mutated to alanine as shown schematically. HEK293T cells were seeded in 6-well plates overnight, and then each well was co-transfected with pCMV4-3HA**-**IκBα (1μg) and either WT or indicated mutant pcDNA6-p62-Myc (1μg) for 48 h. Whole cell lysates (500 μg) were used for co-IP with Myc-trap (50 μL), followed by IB analyses with HA- and Myc-antibodies.

To identify the corresponding IκBα-hotspots involved in p62-interactions, we carried out IκBα-structural deletion analyses (**Fig. 5**). For reference, IκBα-NF-κB interactions under basal conditions are depicted (**Fig. 5A;** 41). Given that the positively charged p62-microregion is a hotspot for IκBα-interaction, we aimed at identifying negatively charged IκBα-residues either adjacent or in the vicinity of its p62-crosslinked K- or C-residues on the IκBα structural surface. As depicted in magenta (**Fig. 5A**), two plausible negatively charged sites were identified: E_85_E_86_ adjacent to K_87_, and E_200_ adjacent to K_238_ (**Fig. S5**). Individual site-directed mutation of each of these E-residues to Ala failed to appreciably alter p62-IκBα-interactions (*Data not shown*). We then designed IκBα-truncation mutants to sequentially delete domains containing these plausible p62-interaction hotspots. In these analyses, plasmid constructs encoding various HA-tagged IκBα-subdomains (depicted in **Fig. 5B**) were co-transfected into HEK293T cells with or without the p62-Myc plasmid (**Fig. 5C**). To assess the extent of p62 protection against IκBα proteasomal degradation, and to maximize the detection of IκBα-levels through stabilization, these cells were also pretreated with or without the proteasomal inhibitor MG-132 (**Fig. 5D**). Cells co-expressing HA-IκBα and p62-Myc were then subjected to anti-HA co-IP to determine the amount of interacting p62 as in **Fig. 1A**. Relative to the WT-protein, IκBα-proteins with deletions of its PEST-subdomain and/or the Ub-independent degradation degron (constructs 1-206 and/or 1-287), were inherently more stable, but p62 co-expression further enhanced their stability, and MG-132 had minimal further effect. IκBα-proteins with substantial deletions of the N-terminal residues (constructs 44-317, 67-317, 104-317), were inherently less stable, but their stability was greatly rescued by p62 co-expression, with minimal further effect by MG132 treatment (**Fig. 5C**). This suggested that p62 could still interact and stabilize IκBα with either the C-terminal AR5, AR6 and PEST domains deleted, or the N-terminal regulatory region and AR1 domains deleted. By contrast, the IκBα-proteins with both N- and C-terminal regions deleted, (constructs 104-199 and/or 104-209), and thus excluding all the potential p62-interaction sites were inherently unstable and p62-co-expression failed to rescue these proteins. Only cell-treatment with MG132 was capable of appreciably stabilizing these 2 truncation mutants. Remarkably, construct 104-199, with E_200_ deleted, totally lacked its p62-mediated stabilization (**Fig. 5C**, *right panel*) and interaction, even in the presence of MG132 (**Fig. 5D**, *lower right, IP/HA data*). Together these findings (**Figs. 4 & 5**) reveal that IκBα-protein stability and molecular p62-IκBα-protein interactions not only rely on both N- and C-terminal IκBα-subdomains, but are also largely dependent on the positively charged p62-residues in its IR-subdomain. Inspection of the existing crystal structure of the IκBα-NF-κB complex (41; **Fig. 5A**) reveals that these p62-interacting IκBα-regions are on the IκBα-interface opposite to that which binds NF-κB (41-43) and thus are not expected to competitively disrupt its NF-κB-association.

**FIGURE 5.**
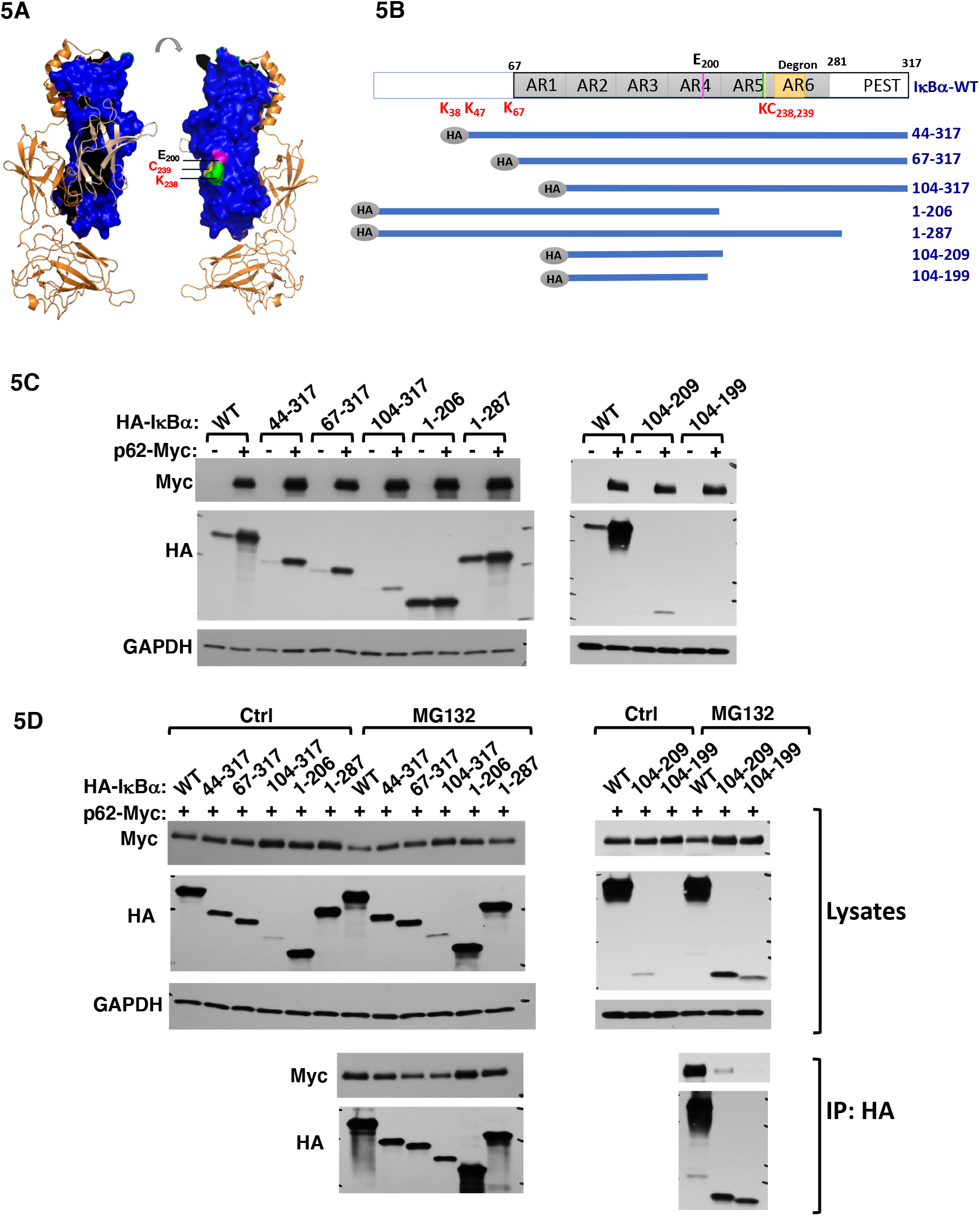
IκBα p62-interaction region verified by structural mutagenesis. **A**. Scheme of a series of N-terminal and C-terminal truncation mutants of IκBα were designed and constructed corresponding to the hotspots identified (**Figs. 4** and **5**). **B**. Crosslinked sites (green) and the negatively charged amino acids adjacent to the crosslinked sites (magenta) were highlighted on the surface of IκBα structure (blue) complexed with p65-p50 (depicted as orange ribbon). **C**. HEK293T cells were seeded in 6-well plates overnight, and then each well was transfected with of pCMV4-3HA**-**IκBα WT or indicated mutants (1μg) with or without pcDNA6-p62-Myc for 48 h. Whole cell lysates (10 μg) were subjected to IB analyses with GAPDH as the loading control. **D**. HEK293T cells were seeded in 6-well plates overnight, and then each well was co-transfected with pCMV4-3HA**-**IκBα WT or indicated mutants (1μg) along with pcDNA6-p62-Myc for 42 h. Cells were treated with vehicle control (0.05% methanol) or MG132 (20 μM) or 6 h. Whole cell lysates (500 μg) were used for co-IP with HA-agarose (50 μL), eluents as well as cell lysate input (10 μg) were used for IB analyses with HA- and Myc-antibodies.

### Influence of p62-IκBα-interactions on intracellular IκBα-protein partnerships as monitored through in-cell APEX-proximity labeling analyses

The collective findings of the sequence-deletion analyses coupled with the finding that some mutant IκBα-proteins (i.e. constructs 104-199 and 104-209) found incapable of significant p62-interaction, could be appreciably stabilized only by MG-132, but not by p62-coexpression (**Fig. 5D**), indicated that efficient p62-interaction affords cellular protection against rapid IκBα-proteolytic turnover and could thus effectively influence its cellular function and partnerships.

To scrutinize these cellular p62-IκBα-protein interactions in greater detail and elucidate additional participants and/or interacting partners if any, we conducted in-cell APEX-proximity labeling analyses. Our aim was to determine the relative differences in the IκBα interactome with or without p62 co-expression. We compared APEX-IκBα fusion labeled biotinylated proteins with or without p62 co-expression, after filtering out the non-specific binders by employing GFP-IκBα co-expressed p62 as a negative control (**Fig. 6A, Fig. S6**).

Not surprisingly, Volcano plot analyses of cells co-expressing both APEX-IκBα and p62-Myc relative to those expressing APEX-IκBα alone revealed that upon p62-coexpression many cellular interactants were either significantly increased or decreased several-fold (**Fig. 6A**). However, consistent with the p62-interaction with residues on the IκBα-interface opposite to that of its NF-κB-interaction, p62-coexpression did not appreciably affect IκBα-associated NF-κB2 (p52) and RelA (p65)-levels, whereas NF-κB1 (p50) and Rel C-levels were only slightly (<2-fold) reduced. By contrast, the levels of several 19S proteasomal regulatory particle ATPase and non-ATPase subunits as well as 20S proteasomal core subunits interacting with IκBα were markedly reduced upon p62-coexpression (**Fig. 6B**), quite consistent with the impaired proteasomal degradation and greater proteolytic stability observed upon its cellular p62-interaction. On the other hand, the interactions of the IκBα-p62 complex with the mitochondrial 28S ribosomal complex tended to increase (**Fig. 6C**), suggesting greater IκBα localization to the mitochondrial matrix, possibly due to enhanced intramitochondrial p62-mediated IκBα-trafficking.

### Disruption of intracellular p62-IκBα protein interactions: Physiological and pathophysiological consequences

Collectively, the above findings revealed that by influencing IκBα proteolytic stability, p62 could regulate the relative robustness of the cellular NF-κB-IκBα association and consequently NF-κB-activation (**Fig. 7**). To probe any such potential p62 regulation, we first examined TNFα-elicited NF-κB-activation and subsequent feedback termination in WT and p62^-/-^-MEF cells (**Fig. 8A**). In WT MEF cells, TNFα-treatment elicited rapid destruction of the cytoplasmic NF-κB-associated IκBα, followed by unleashing of NF-κB, its nuclear translocation (as evidenced by the increased nuclear p65 content) and its activation [reflected by the increased nuclear phosphorylated p65 (P-p65) content]. The consequent transcriptional activation of the NF-κB-target IκBα-gene resulted in rapid *de novo* IκBα protein synthesis that effectively terminated the TNFα-elicited NF-κB activation (evidenced by decreased nuclear p65 and P-p65 content), consistent with a classical NF-κB-IκBα negative feedback loop (9, 11). By contrast, in p62^-/-^-MEF cells, the newly synthesized IκBα pool being much more labile in the absence of p62, was unable to vigorously compete for the DNA-associated NF-κB and escort it out of the nucleus, thus disrupting the NF-κB-IκBα negative feedback loop. As a result, NF-κB persisted in the nucleus and its activation was greatly prolonged (**Fig. 8A**).

**FIGURE 6.**
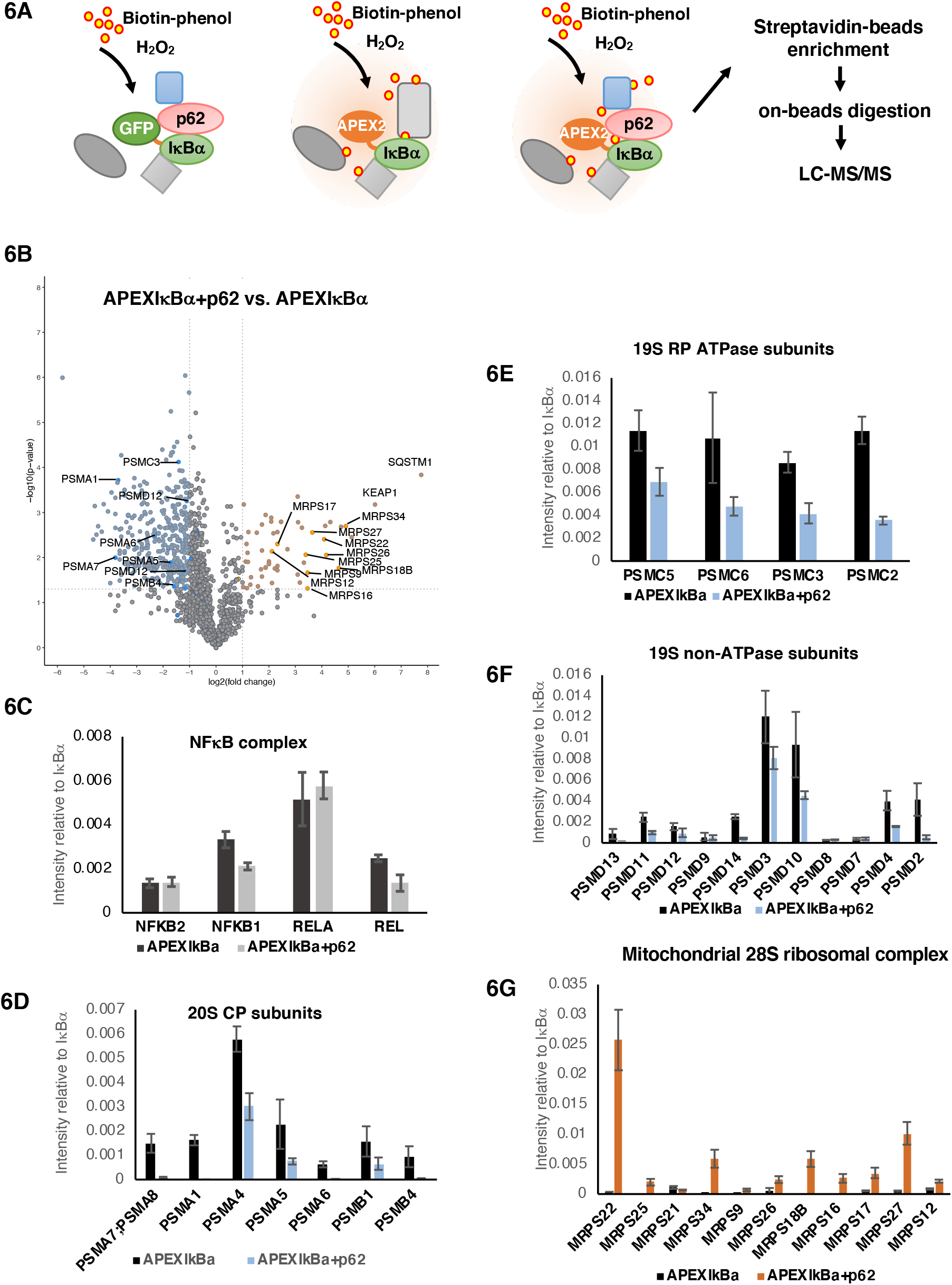
APEX captures proteins that differentially interact with IκBα upon p62 co-expression. **A**. Scheme of APEX workflow to identify and compare proteins that specifically interact with IκBα when it was expressed alone, or when it was co-expressed with p62. GFP-IκBα was used as a negative control for APEX-labeling. **B**. Volcano plot of relative protein spectral count derived from APEX-IκBα and APEX-IκBα + p62 proteomic analyses. Data from 4 independent experiments are presented as the mean. Proteins found to interact to a greater extent with IκBα when p62 is present are highlighted in orange (Fold change > 2, p value < 0.05), proteins found to interact to a greater extent with IκBα when p62 is absent are highlighted in blue (Fold change > 2, p value < 0.05). **C**. Bar chart of the relative intensities of identified NFκB complex proteins derived from APEX-IκBα and APEX-IκBα + p62 proteomic analyses. Data presented as mean ± SD (n = 4). **D**. Bar chart of the relative intensities of identified 20S core proteasomal subunit proteins derived from APEX-IκBα and APEX-IκBα + p62 proteomic analyses. Data presented as mean ± SD (n = 4). **E**. Bar chart of the relative intensities of identified 19S proteasomal regulatory particle ATPase subunit proteins derived from APEX-IκBα and APEX-IκBα + p62 proteomic analyses. Data presented as mean ± SD (n = 4). **F**. Bar chart of the relative intensities of identified 19S regulatory particle non-ATPase subunit proteins derived from APEX-IκBα and APEX-IκBα + p62 proteomic analyses. Data presented as mean ± SD (n = 4). **G**. Bar chart of the relative intensities of identified mitochondrial 28S ribosomal complex proteins derived from APEX-IκBα and APEX-IκBα + p62 proteomic analyses. Data presented as mean ± SD.

**FIGURE 7.**
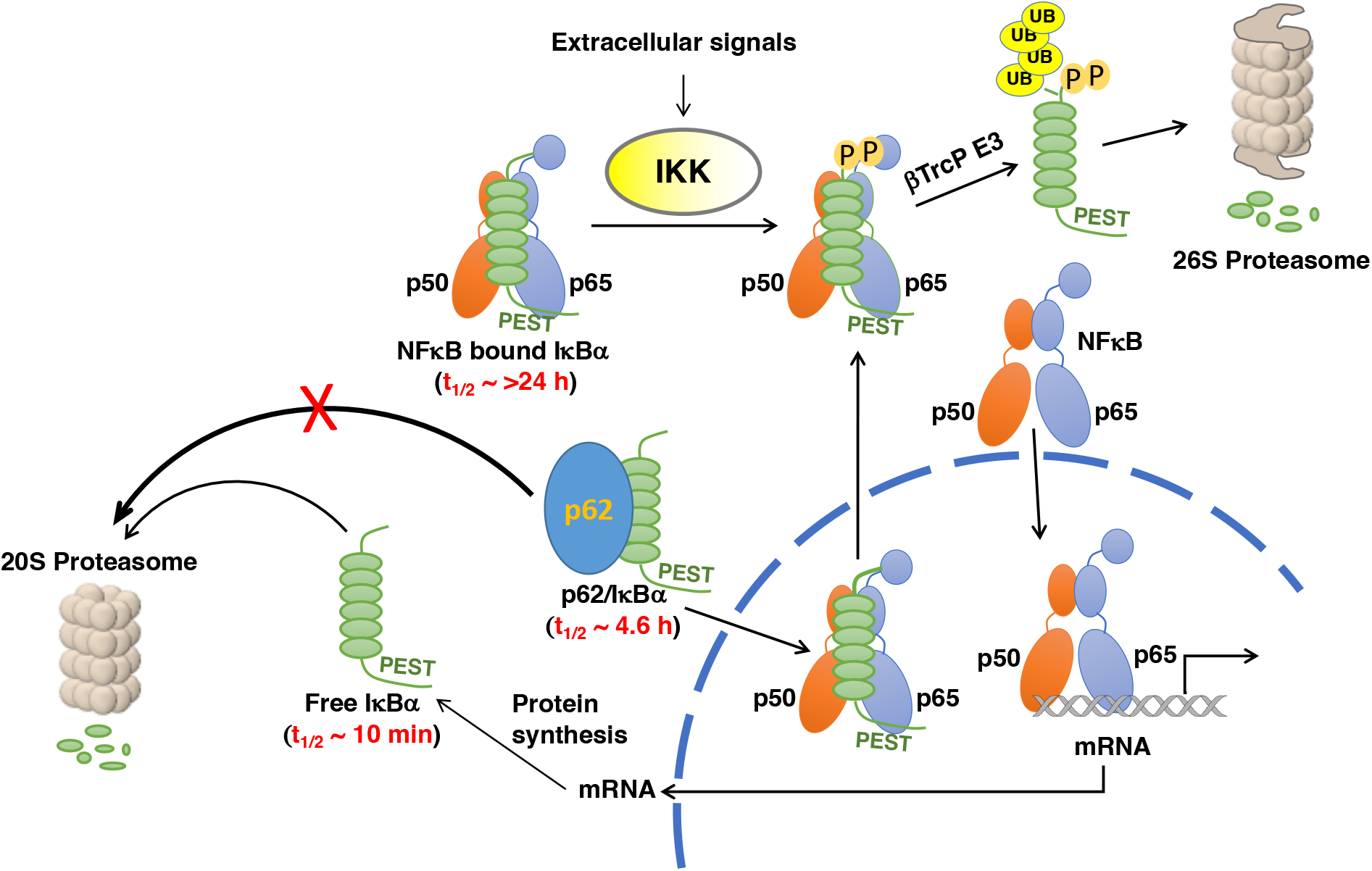
A scheme of the cellular pathway of NF-κB activation and IκBα-mediated feedback loop, indicating a potential role of p62 in IκBα-stabilization and -feedback. Because p62 co-expression also impairs IκBα-association with 26S cap subunits, p62-association may also block its 26S proteasomal degradation

**FIGURE 8.**
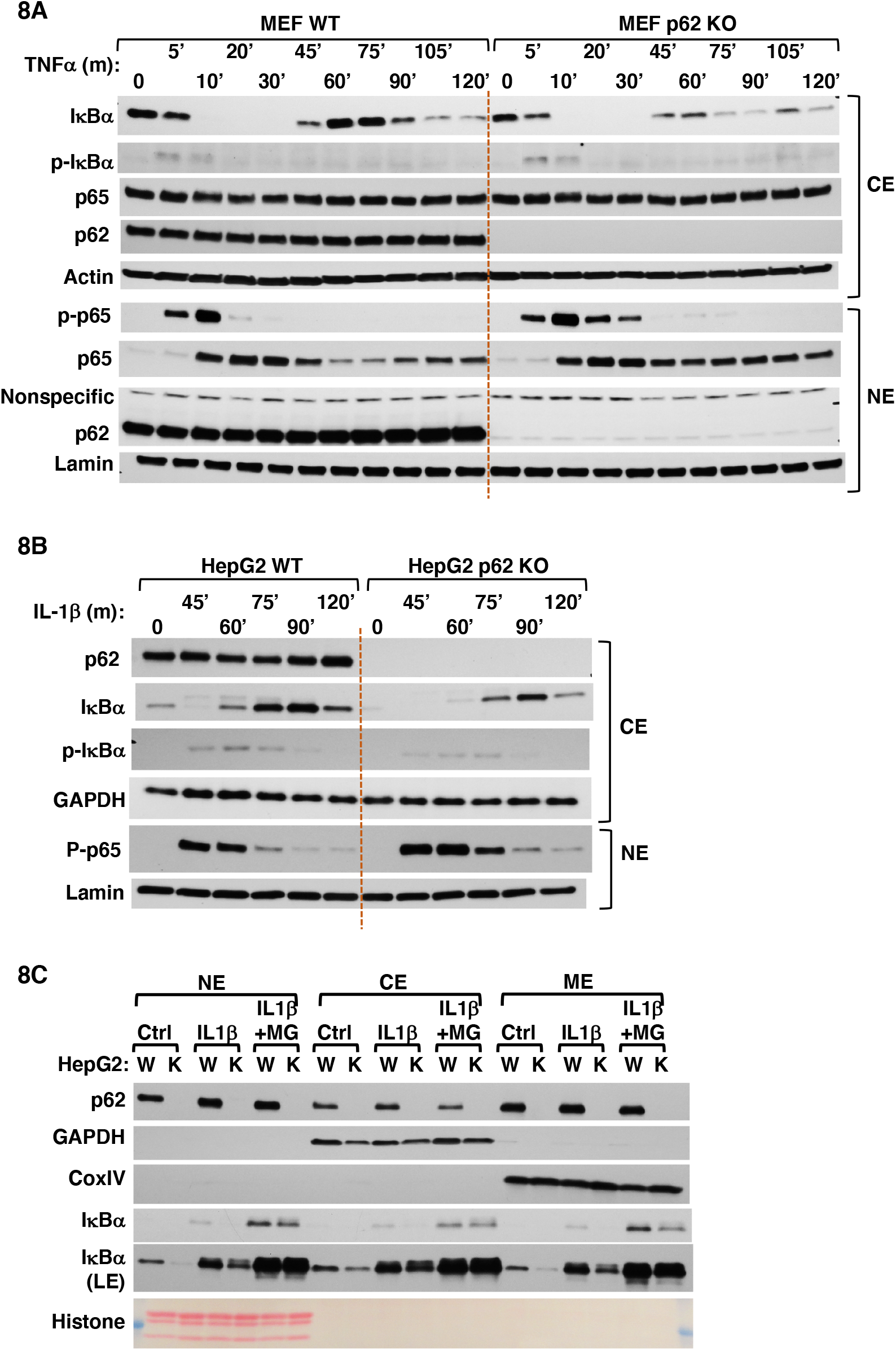
p62-KO resulted in more intense and prolonged NF-κB response. **A**. p62 WT and KO MEF-cells were stimulated with TNFα (20 ng/mL). Cells were collected at indicated times and fractionated into cytoplasmic (CE) and nuclear (NE) extracts. Extracts (10 μg) were used for IB analyses. **B**. p62 WT and CRISPR-KO HepG2 cells were pulse stimulated with IL-1β (20 ng/mL) for 15 min. Cells were collected at indicated times and fractionated into cytoplasmic (CE) and nuclear (NE) extracts. Extracts (10 μg) were used for IB analyses. **C**. p62 WT (W) and CRISPR KO (K) HepG2 cells were either untreated (Ctrl) or pulse-stimulated with IL-1β (20 ng/mL) for 15 min, then incubated with or without MG132 (MG; 20 μM) for an additional 75 min. Cells were collected and then separated into nuclear (NE), cytoplasmic (CE), and mitochondria (ME) extracts. Extracts (10 μg) were used for IB analyses with GAPDH as the loading control as well as a cytoplasmic marker, histone H3 as the loading control and nuclear marker, and CoxIV as loading control and mitochondrial marker.

Similar disruption of the NF-κB-IκBα negative feedback loop was also observed in p62^-/-^-HepG2 cells (**Fig. 8B**) upon stimulation with IL-1β, another inflammatory cytokine that is a more potent NF-κB transcriptional activator than TNFα in HepG2-cells and mouse liver hepatocytes (**Fig. S7A**). In these IL-1β-stimulated p62^-/-^-HepG2-cells, the lack of p62-mediated stabilization caused greater proteolytic degradation of the newly synthesized IκBα, which could be prevented by MG132 treatment (**Fig. 8C**, *cytosolic extract (CE) panel*). As a result, the nuclear NF-κB activation (P-p65) and persistence was prolonged relative to those in the corresponding p62^+/+^-WT cells (**Figs. 8B; S7B**), establishing that p62 was indeed required for a robust NF-κB-IκBα negative feedback loop response.

To further verify the physiological and pathophysiological relevance of intact p62-IκBα-interactions *in vivo*, we generated a mutant mouse upon liver-specific genetic ablation of the p62 residues 68-252 (p62mut). This p62mut thus lacks the critical hepatic IκBα-p62-interacting region (IR, R_183_, R_186_, K_187_ and K_189_) (**Fig. 9A**). After IL-1β stimulation of p62mut mouse hepatocytes with disrupted p62-IκBα-interactions (**Fig. 9B**), a similarly reduced level of newly synthesized IκBα along with a prolonged nuclear NF-κB (P-p65) activation was observed as in p62^-/-^-HepG2 cells (**Fig. 8B**). Liver histological analyses revealed that while the WT mice exhibited an age-dependent mild hepatic inflammation, this was considerably further exacerbated in the p62mut mice, effectively leading to their increased mortality by 16 months of age (**Fig. 9C**). Such age-dependent relative increases in liver inflammation in p62mut-mice were associated with corresponding increases in their hepatic levels of inflammatory markers [IL-6, IL-1β, TNFα and SPP-1 (osteopontin/secreted phosphoprotein 1)] over age-matched WT controls (**Fig. S8**). These findings conclusively established that disruption of physiological p62-IκBα-interactions has severe pathophysiological consequences.

**FIGURE 9.**
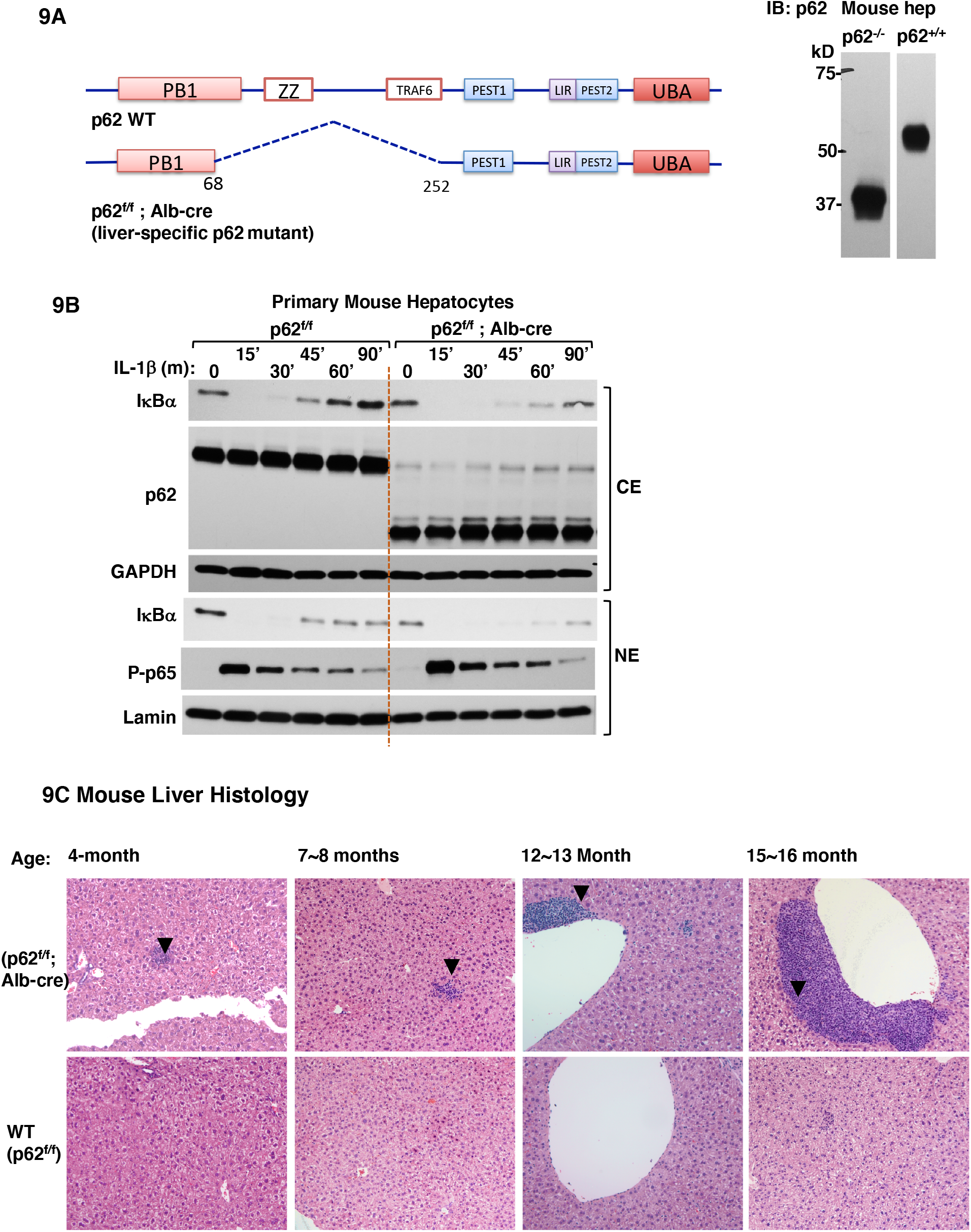
Transgenic mice with a liver-specific p62 deletion of the IκBα-interacting region (p62mut) exhibited increased inflammation upon aging. **A**. A scheme indicating the p62-region deleted in the liver-specific p62mut mice, and p62 Western immunoblot of hepatocytes isolated from p62 WT and p62mut mice. **B**. Primary hepatocytes from these WT and p62mut mice were cultured for 5 d, and then pulse stimulated with IL-1β (20 ng/mL) for 15 min and then harvested at indicated times and fractionated into cytoplasmic (CE) and nuclear (NE) extracts. Extracts (10 μg) were used for IB analyses. **C**. Mice at indicated age ranges (4-16 months) were sacrificed and liver pieces were used for H&E stain and histology. Arrows indicate inflammatory regions.

## DISCUSSION

Collectively, our findings detailed above reveal an intimate intracellular p62-IκBα association, which stabilizes cellular IκBα levels by limiting its interaction with the 20S/26S proteasome and subsequent degradation, thereby greatly extending its life-span. The enhancement of such interactions upon cell-/lysate-pretreatment with NEM which would abrogate SH-elicited post-translational modifications such as ubiquitination and/or sumoylation, suggests that p62 preferentially interacts with the native, unmodified IκBα, thus possibly favoring the nascent, *de novo* synthesized protein.

Our characterization of such p62-IκBα-interactions through various complementary approaches such as structural deletion, in-cell CLMS and/or site-directed mutagenesis analyses of each protein as well as cell transfection/co-immunoprecipitation assays have revealed structural hotspots critical to this protein-protein interaction (**Fig. 10**). Thus, in p62 the basic residues (R_184_, R_186_, K_187_ and K_189_) appear critical for its IκBα-association, as this association is nearly abrogated upon their Ala-mutation. Intriguingly, this p62-site coincides with its designated NLS1 (residues 183-194) (38). Although chemical-crosslinking also revealed IκBα-interactions with p62 K_13_, Ala-mutation of this residue failed to appreciably affect p62-IκBα-interactions. Similar IκBα-analyses identified two major hotspots: One in its N-terminus centered around residues 38-67, and the other around residues 238/239 in its 5^th^ AR-domain (ARD). The sufficiently close proximity of the latter p62-crosslinked site to its Ub-independent degron (residues 243-280) in its 6^th^ ARD (32), which is normally masked through NF-κB-binding, most likely accounts for the IκBα proteolytic stability conferred by both p62- as well as NF-κB-association. However, following proinflammatory stimulation of nuclear NF-κB-activation, this degron in *de novo* synthesized IκBα-protein would be unshielded, and subject to rapid proteasomal degradation. Strategically, cytoplasmic association with p62, which is also concurrently induced upon NF-κB-activation (13-16), would enable IκBα survival long enough to insure its nuclear import and consequent tight regulation of the relative duration of NF-κB-mediated transcriptional activation cycle. Consistently, in the absence of cellular p62, the proteolytic instability of *de novo* synthesized IκBα resulted not only in prolonged activation of NF-κB (P-p65), but also in its nuclear persistence beyond the normal TNFα- or IL-1β-elicited IκBα-NF-κB feedback cycle (**Fig. 8**). Our findings would thus argue that in addition to the two cellular IκBα-pools known to exist i.e. a major (≈ 85%), long-lived cytoplasmic NF-κB-complexed IκBα-pool (t_1/2_ ≈ days), and a minor (≈ 15%) much shorter-lived free IκBα-pool (t_1/2_ ≈ 10-15 min) (31, 44), a third cellular pool of p62-complexed IκBα (t_1/2_ of ≈ 4.6 h) must exist to shield *de novo* synthesized “free” IκBα from rapid proteasomal degradation, thereby enabling its nuclear import and subsequent effective negative feedback regulation of nuclear NF-κB activation. Given that the p62-interaction occurs with the IκBα-interface directly opposite to that of its NF-κB-binding (41-43), the possibility of p62-piggy backing with the longer lived IκBα-NF-κB pool is in principle conceivable. However, this is precluded by the fact that no NF-κB was ever detected in the DSS- or SIAB-crosslinked IκBα-p62 complexes upon our proteomic verification. More importantly, such a close p62 association with the newly synthesized IκBα would circumvent the rapid proteasomal degradation that free, nascent IκBα with its readily accessible PEST and Ub-independent and -dependent degrons could otherwise be subject to (31, 32, 44). This important protective chaperone role notwithstanding, the strategic inflammatory stimuli-elicited NF-κB-mediated concurrent transcriptional activation of both p62 and IκBα, and the above documented association of these two *de novo* synthesized proteins, leads us to propose that this association may additionally enable efficient nuclear import of IκBα required to terminate the NF-κB-activation cycle.

**FIGURE 10.**
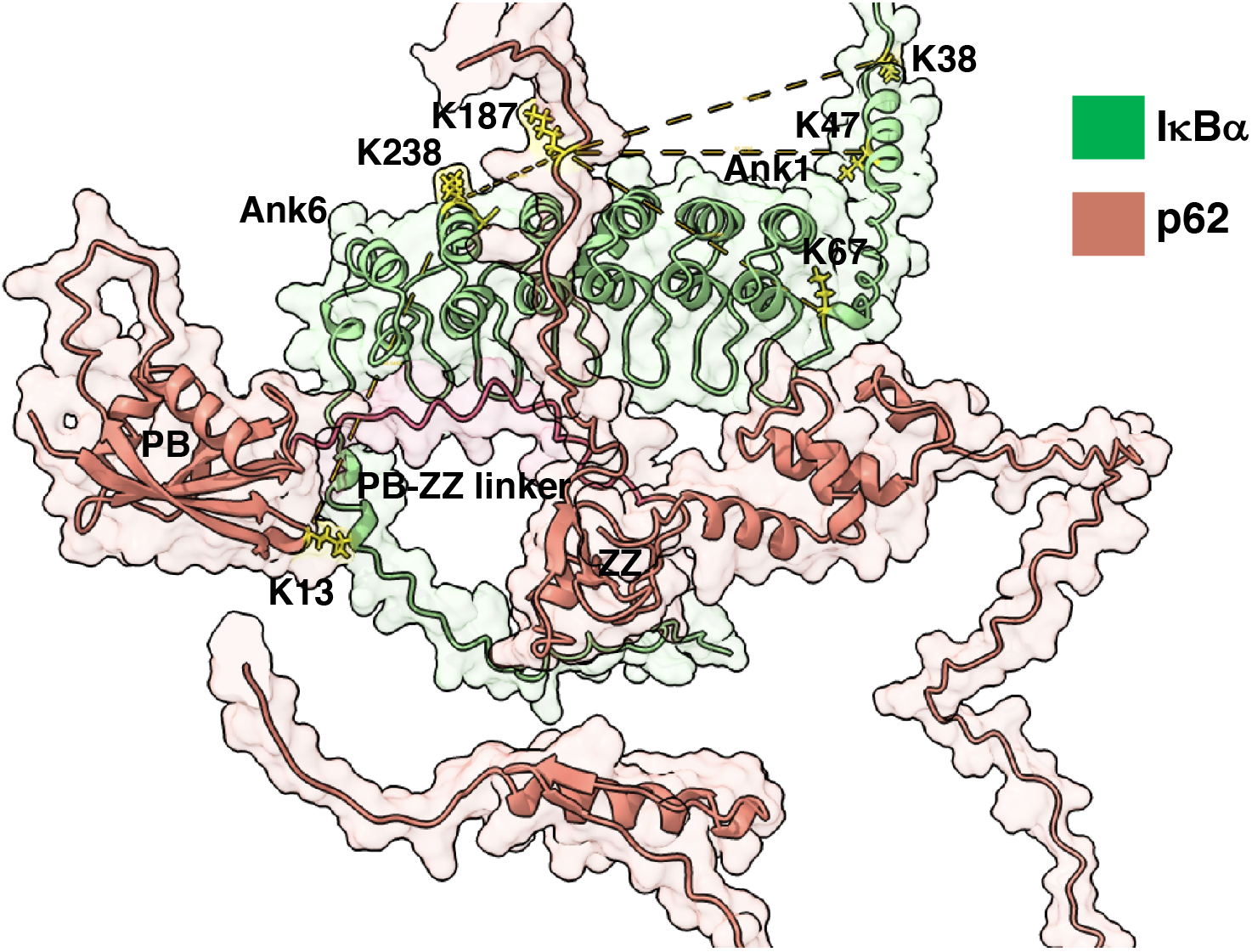
AlphaFold predicts an interaction between the p62 PB-ZZ linker and IκBα ARs. The p62-IκBα dimer interaction was modeled using AlphaFold v2.1.2. All five of the independent modeling runs predicted similar structures with contacts between the PP-ZZ linker of p62 and 4-6 ARs of IκBα with Predicted Aligned Errors (PAE) values between 15-20Å (**Fig. S9**). The top ranked model is shown with the PB-ZZ linker (residues 102-123 shown in magenta). Crosslinked Lys-residues are shown in yellow. K187 on p62 occurs on a flexible loop that is poorly localized in the AlphaFold model. K13 on p62 lies in the PB-1 domain which mediates oligomerization between p62 protomers. Crosslinks between K13 and C239 are not plausible as depicted and most likely originate from a p62 oligomer-IκBα interaction.

IκBα nuclear import given its small molecular size and lack of canonical structural NLS, was once believed to occur via simple diffusion (9, 45). However, this view was modified when a novel class of non-canonical cis-acting discrete nuclear import sequences (NIS) that consist of a hydrophobic residue cluster within the IκBα-N-terminal 114-124 residues in its 2^nd^ ARD was identified as the predominant, albeit not sole NIS (46). This 2^nd^ ARD hydrophobic cluster is highly conserved among other IκB family proteins, and all the IκB-proteins commonly sharing this 2^nd^ ARD-hydrophobic cluster, exhibit the apparently conserved feature of nuclear import (46). Furthermore, insertion of this ARD hydrophobic cluster into other NLS-deficient proteins confers nuclear import capabilities on them (46). Although such a nuclear import does not require Rel-proteins, this ARD hydrophobic cluster is critical both for its p65-association and blocking its DNA-binding, as it is masked upon cytoplasmic p65-binding (44, 46). A fully functional AR-derived IκBα-NIS harboring such an ARD hydrophobic cluster was subsequently proposed to consist of N-terminal β-hairpin, 2 α-helices and N-terminal β-hairpin of the adjacent ARD (46). However, whether, this discrete NIS functioned autonomously, or required a piggy-back mechanism and/or another receptor that was importin-assisted remained unclear. Turpin et al. (47) similarly concluded that IκBα nuclear import was not via passive diffusion but via a specific active cytosol/energy-dependent process whose *in vitro* reconstitution required IκBα ARDs, Ran-GDP, α- and β-importin and an as yet unknown cytosolic factor(s). Because passage of the cytosolic fraction through an affinity GST-IκBα[68-243] column but not through a GST-column aborted the capacity of the cytosolic fraction to fully reconstitute the nuclear IκBα import process, they argued that ARDs contained in the IκBα[68-243] residue domain could sequester and thus effectively deplete this cytosolic factor(s) (47). By contrast, this depleted cytosolic fraction could fully support the nuclear import of a BSA-NLS-FITC protein containing a basic amino acid stretch NLS (47). This led to their proposal that most likely, IκBα nuclear import machinery additionally included an unknown cytosolic factor containing a basic NLS for piggy-backing IκBα (47). Subsequently, Lu et al. (48) further refined this concept, and proposed a novel, importin-*in*dependent RanGDP/AR nuclear import pathway that relied on an IκBα nuclear import code: 13^th^ hydrophobic residues of two consecutive ARs that interact with RanGDP to chaperone IκBα through the nuclear pore complex (NPC) aided by its specific interactions with the FG-rich NPC-components, Nup153 and RanBP2 (49-53). These studies were largely conducted following co-transfections of these FITC-tagged AR-derived constructs into digitonin-permeabilized cells and subsequent pull-downs with GST-RanGDP (48). However, to our knowledge, intact IκBα has never been actually documented to similarly localize to the nucleus solely via this nuclear import pathway. Our previous proteomic findings of ZnPP-aggregates, revealing the existence of a very close and consistently strong association between IκBα, Nup153 and RanBP2, but not with either importin α or β, also led us to conclude that RanGDP due to its reported affinity for Nup153 and RanBP2 (49-53) was most likely involved in IκBα nuclear import (17).

However, the IκBα[68-243] domain that was suggested to sequester and thus deplete the critical cytosolic factor required for IκBα nuclear import (47), not only contains its ARDs, but also regions adjacent to K_67_ as well as K_238_/C_239_, residues we found situated in its p62-interacting hotspots. Thus, p62 could very well qualify as one, if not, the only depleted cytosolic factor. This consideration coupled with our findings described above, make it plausible that p62 could represent a *bona fide* piggy-back for IκBα nuclear import. Several intrinsic features uniquely qualify p62 for such a role: (i) p62 reportedly shuttles continuously in and out of the nucleus (38). Although its NLS1 would be masked by IκBα-association, its structurally more dominant basic NLS2 (K_264_RSR_267_-residues) (38), being eminently accessible, would enable participation in such nuclear import. (ii) Its NF-κB-elicited concurrent transcriptional induction with IκBα (13-16) could readily promote the association of these two *de novo* synthesized proteins. (iii) The intimate cellular association of the two proteins, and the observed considerably reduced nuclear levels of IκBα with correspondingly increased NF-κB nuclear preponderance and persistence upon p62-KO (**Fig. 8**). Intriguingly, in IL-1β-stimulated HepG2 cells, even though upon MG132-mediated proteasomal inhibition, cytoplasmic IκBα levels in p62-KO cells could be restored to the same level as in WT, the nuclear IκBα levels were consistently reduced in p62-KO cells relative to corresponding WT controls. This additionally argues that p62 may not solely protect IκBα against proteasomal degradation, but also function in its nuclear import. (iv) p62 interaction with the IκBα-interface opposite to that of its NF-κB-interaction in such p62-IκBα complexes (41-43), far from impeding, would strategically favor IκBα-hand-off for normal interactions with the DNA-bound NF-κB, required for the termination of each NF-κB-activation cycle through IκBα-mediated DNA-dissociation and subsequent escort out of the nucleus.

In addition to providing the much sought after “piggy-back” for IκBα nuclear import, it is plausible that p62 may similarly function in mitochondrial import of IκBα. The p62-mediated negative regulation of inflammation through mitophagy has been previously documented in macrophages (54). Our findings reveal that p62 co-expression led to increased IκBα localization to the mitochondria matrix (**Fig. 6G**); consistently, very little IκBα is found in the mitochondrial extracts of p62-KO HepG2 cells relative to those of corresponding p62-WT cells, both under basal conditions and upon IL-1β-stimulation (**Fig. 8C**). And although, MG-132 rescues some of this nuclear IκBα content, it is still appreciably lower than that found in corresponding IL-1β-treated p62-WT cells. Furthermore, our findings of enhanced association of mitochondrial 28S ribosomal complex proteins with IκBα in the presence of p62-coexpression relative to its absence, also support such a p62-assisted mitochondrial IκBα-import (**Fig. 6G**) Given that p62 is documented to promote proliferation and reduce apoptosis (55, 56) and mitochondrial IκBα is thought to serve a unique protective role against apoptosis, it is conceivable that by enhancing IκBα-mitochondrial import, p62 may similarly play a beneficial role in hepatic inflammatory responses.

Collectively, our findings document that p62 interacts with IκBα through a novel protein-interaction region which is independent of its many previously characterized structural protein-interacting regions including that with the innate defense regulator (IDR-1) peptide (34-40, 57). This p62-IκBα-interaction region, however not only comprises its NLS1 (38), but is also precisely the region known to interact with the mutant Cu-Zn superoxide dismutase, SOD1, a mutant linked to amyotrophic lateral sclerosis (ALS) (58). The abrogation of such p62-IκBα interactions upon structural deletion of p62 PB1-domain, reveal that such interactions additionally require p62-oligomerization, as in the case of SOD1 (58). Such IκBα-p62 interactions we have documented are important not only for extending the cellular survival of IκBα (particularly, its readily degradable “free” *de novo* synthesized species), but also may enable its nuclear and possibly mitochondrial import. In HepG2-cells, p62-KO not only enhanced TNFα- or IL-1β-elicited nuclear NF-κB-activation, but also prolonged its nuclear persistence. Furthermore, we found that mouse hepatocytes genetically deficient in this IκBα-interacting p62-subdomain, also exhibit prolonged IL-1β-elicited NF-κB activation. More importantly, our findings that progressively severe hepatic inflammation is observed with aging in intact mice with a genetic deficiency in this hepatic IκBα-interacting p62-subdomain (**Fig. 9C**), reveal that such an IκBα-p62 interaction is physiologically and pathophysiologically relevant. The p62-protein scaffold is known to modulate NF-κB-activation both positively and negatively through direct and/or indirect interactions with various cell-specific interactors and/or signaling effectors at one of its many defined structural subdomains (15, 59). Our findings detailed above further expand this growing repertoire of p62-elicited NF-κB-modulation by revealing a novel, albeit fundamental mode of p62-mediated regulation of NF-κB-activation through its interplay with IκBα. Furthermore, they provide a belated rationale for the then “puzzling” report that in contrast to other scenarios, p62 accumulation in autophagy-deficient iBMK cells led to the inhibition of the canonical NF-κB activation pathway (59, 60).

## ABBREVIATIONS

The abbreviations used are as follows:

ABC: ammonium bicarbonate
AGC: automatic gain control
AR: Ankyrin repeat
ARD: AR-domain
APEX: a 27 kDa engineered monomeric peroxidase (APEX2)
BCA: bicinchoninic acid
BFDR: Bayesian False Discovery Rate
CE: cytoplasmic Extract
CSL: Cell Lysis
CLMS: chemical crosslinking mass spectrometry
Co-IP: co-immunoprecipitation
CHX: cycloheximide
DMEM: Dulbecco’s Modified Eagle high glucose medium
DSS: disuccinimidyl suberate
EThcD: electron transfer/high-energy collision dissociation
FBS: fetal bovine serum
FDR: false discovery rates;
HA: hemagglutinin
HA-IκBα: HA-tagged IκBα
HCD: High collision dissociation
iBMK: immortalized baby mouse kidney cells
IB: Immunoblotting
IL-1β: interleukin-1β
IDR-1: innate defense regulator
IR: intervening region
Lys-C: lysylendopeptidase C
MEFs: mouse embryo fibroblasts
MEM: minimal Eagle’s medium
NCE: normalized collision energy
NE: nuclear extract
NIS: nuclear import sequences
NLS: nuclear localization signal
NPC: nuclear pore complex
NSAF: normalized spectral abundance factor
Nup153: nucleoporin 153
p62 flp/flp: p62-floxed mouse
p62-Myc: Myc-tagged p62
p62mut: p62 genetic mutant mouse
PB-1: Phox and Bem1p-domain
P-p65: phosphorylated p65
RanBP2: a SUMO E3-ligase/Nup358
Ran-GDP: an abundant GTPase involved in nuclear import
SA: streptavidin
SOD1: Cu-Zn superoxide dismutase
SIAB [succinimidyl (4- iodoacetyl)aminobenzoate); SQSTM-1: Sequestosome 1
TNFα: tumor necrosis factor α
TB: TRAF6-binding
Ub: ubiquitin
UPD: Ub-dependent 26S proteasomal degradation
ZnPP: Znprotoporphyrin IX
ZZ: Zn-finger binding motifs

## DATA AVAILABILITY

RAW mass spectrometry data are deposited in the MassIVE repository (https://massive.ucsd.edu) with accession number: **MSV000090324**. (*reviewer password: ikba2022*).

Annotated peaklists supporting the crosslinked spectral assignments are available on MS-Viewer (https://msviewer.ucsf.edu/cgi-bin/msform.cgi?form=msviewer).

Search key (DSS crosslinks): MS-Viewer, ex4ngmrtrj

Search key (SIAB crosslinks): MS-Viewer, l2srapmex0

## ACKNOWLEDGMENTS

We gratefully acknowledge Mr. Chris Her, UCSF Liver Cell & Tissue Biology Core Facility for hepatocyte isolation, and the UCSF Liver Center Pathology Core for the histopathological analyses, both supported by NIDDK Grant P30DK26743. We also sincerely thank Prof. Haining Zhu, University of Kentucky, for providing p62-KO MEF cells generated in Prof. M. Komatsu’s lab, Niigata University, Japan.

## Author Contributions

Y.L., M. J. T. and M.A.C designed the studies and wrote the manuscript. M.A.C supervised the project. Y.L. conducted most of the experiments with MS support, data analyses and interpretation from M. J. T. L. H. carried out the qRT-PCR analyses (**Fig. S8**). A.L.B participated in the discussion of potential MS proximity-labeling approaches to be employed. All authors critically reviewed the manuscript.

