## Supplemental Figures for "In-cell chemical crosslinking identifies hotspots for p62-IκBα interaction that underscore a critical role of p62 in limiting NF-κB activation through IκBα-stabilization"

**S1**

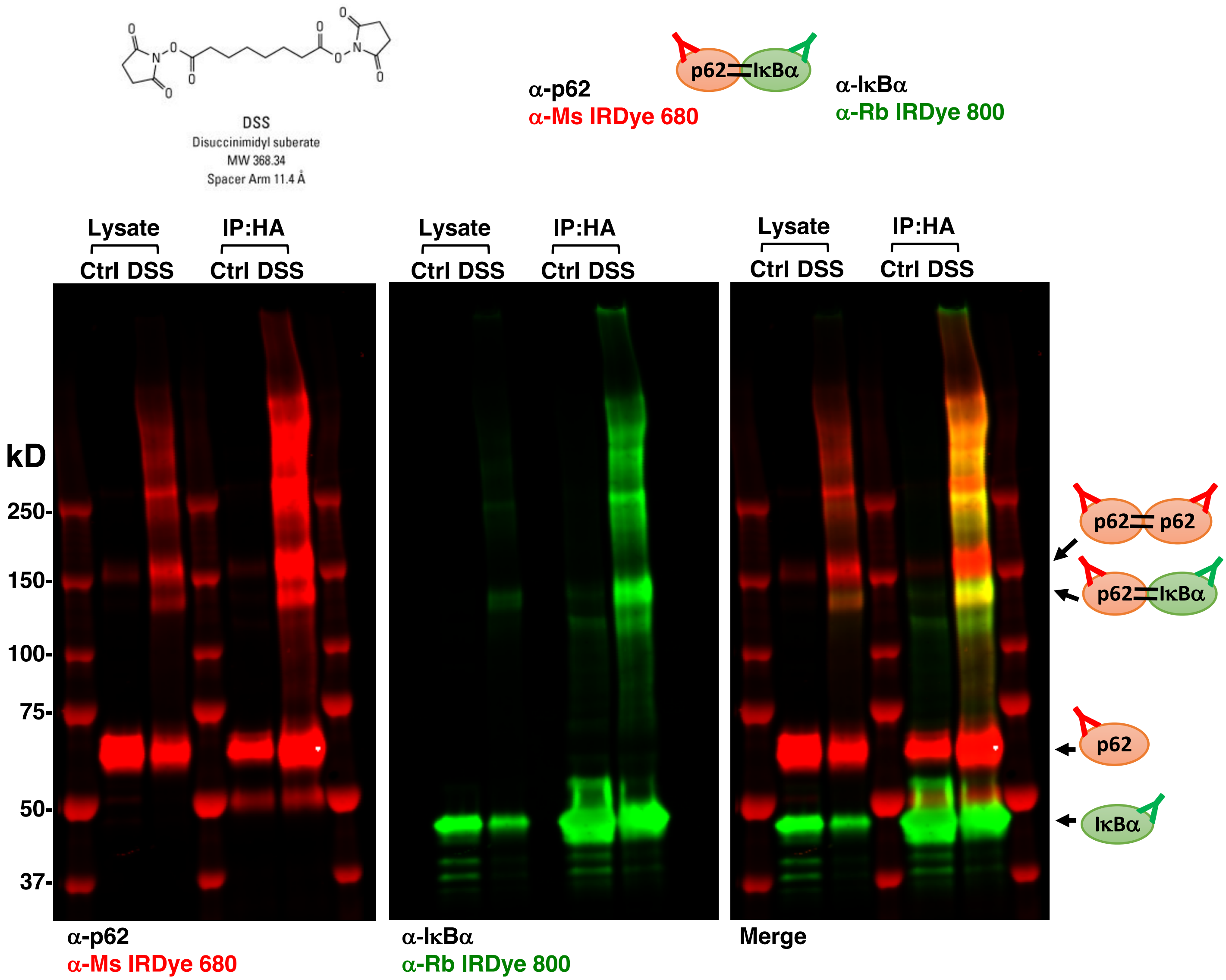


***FIGURE S1. In-Cell chemical crosslinking shows that p62 and IκBα directly interact with each other***. HEK cells overexpressing HA-IκBα and p62-Myc were treated with 5 mM DSS *in vivo*, lysed in RIPA, and then immunoprecipitated with HA-agarose.

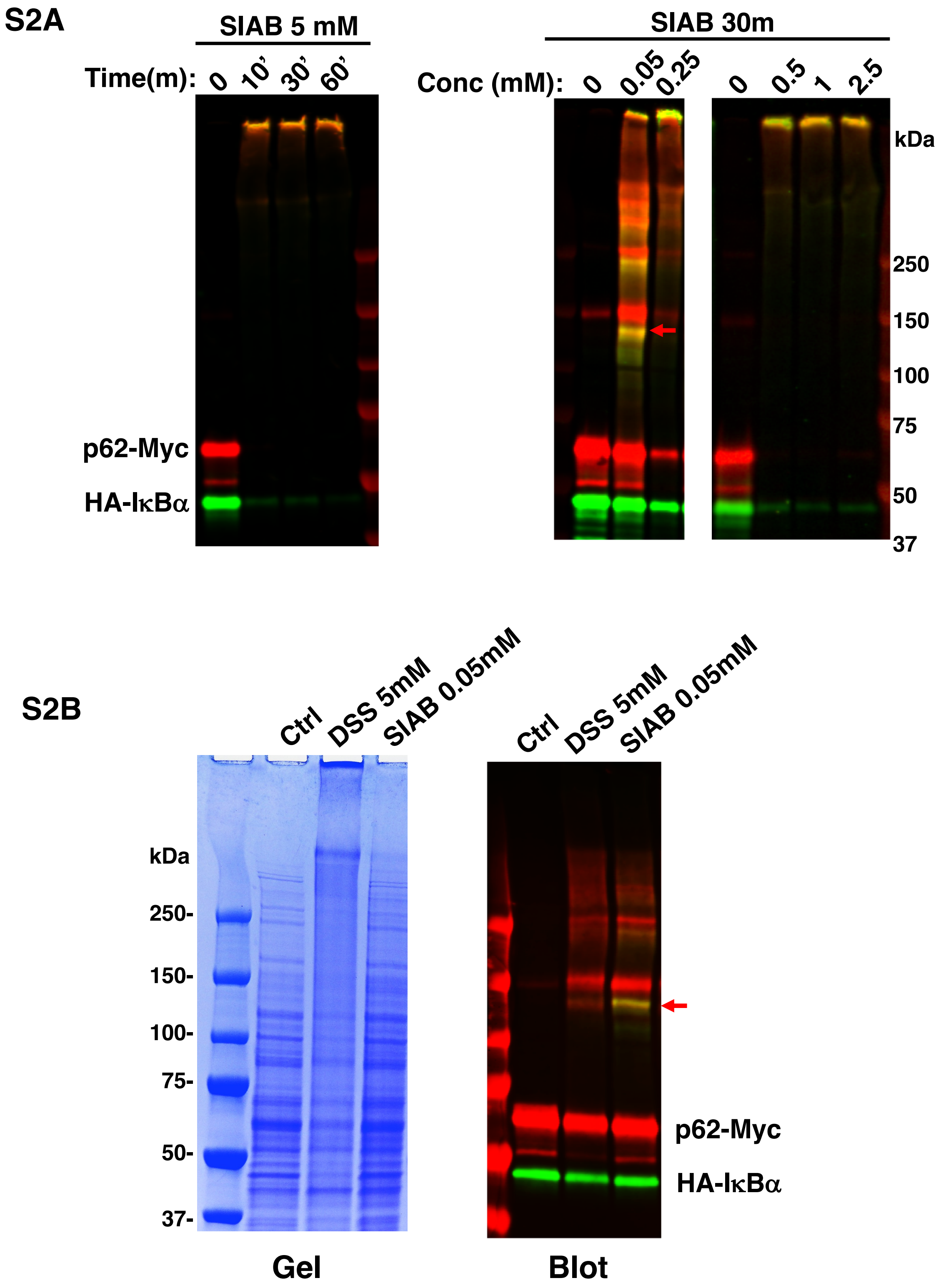


***FIGURE S2. SIAB is more specific and potent in in-cell crosslinking p62 and IκBα*. A.** HEK cells overexpressing HA-IκBα and p62-Myc were treated with SIAB *in vivo* at the indicated concentration and times, lysed in RIPA, and then immunoblotted. **B.** HEK cells overexpressing HA-IκBα and p62-Myc were treated at the indicated concentrations of DSS or SIAB *in vivo,* lysed in RIPA, and then subjeced to SDS-PAGE and immuno-blotting.

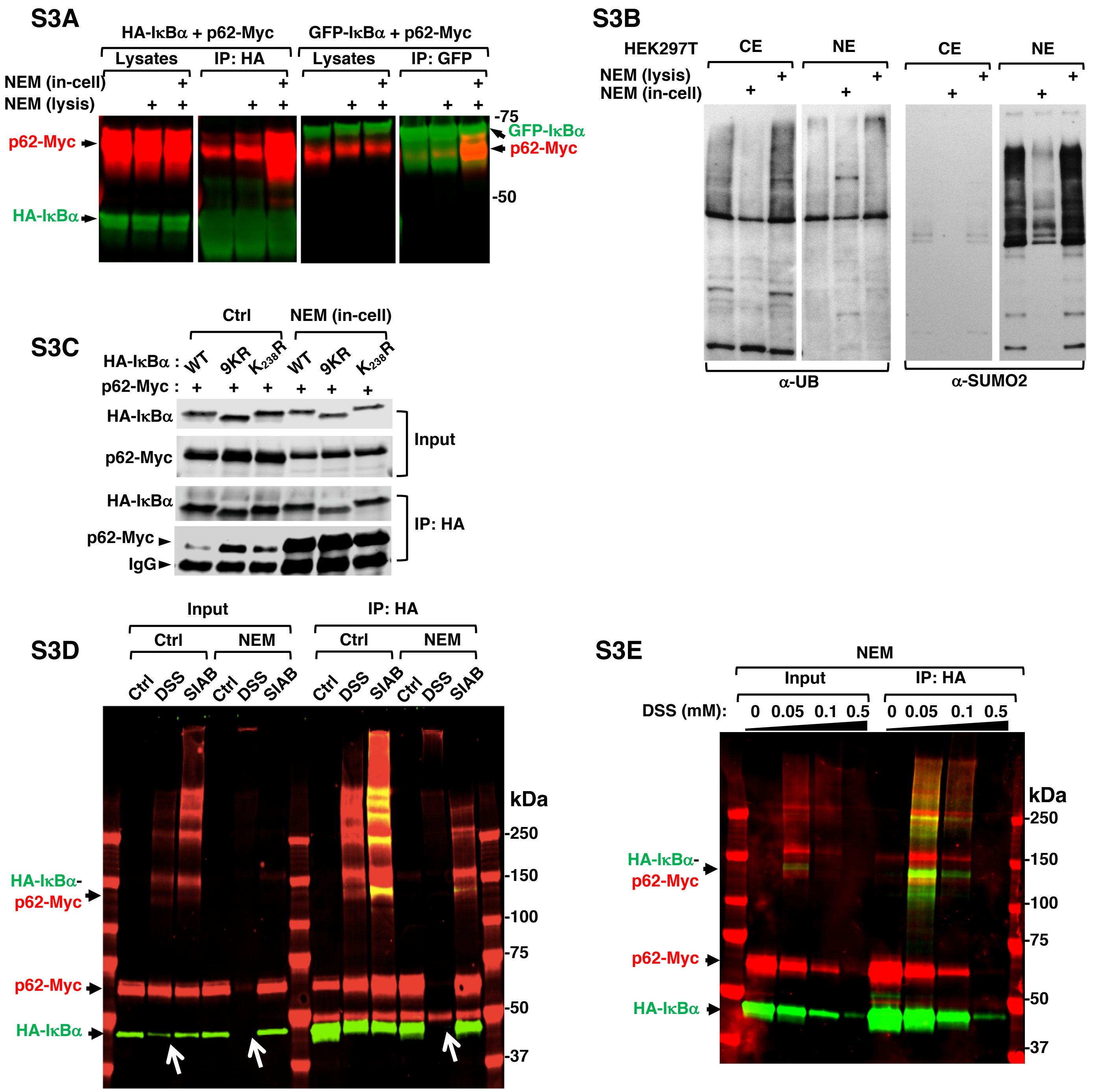


***FIGURE S3. NEM treatment greatly enhanced p62-IκBα interaction and DSS-crosslinking, possibly through inhibiting IκBα PTM. A.*** HEK cells were co-transfected with HA-IκBα and p62-Myc or GFP-IκBα and p62-myc. NEM (10 μM) was added to the RIPA lysis buffer (lysis) or added to live cells 5 min before harvesting (in-cell). Cells were then lysed in RIPA buffer and subjected to immunoprecipitation (IP) with HA-agarose or GFP-trap followed by immunoblotting. **B**. HEK cells were co-transfected with HA-IκBα and p62-Myc. NEM (10 μM) was added to the lysis buffer (lysis) or added to live cells 5 min before harvesting (in-cell). Cells were then fractionated into nuclear extract (NE) and cytosol extract (NE), and subjected to immunoblotting with ubiquitin and SUMO2 antibodies. **C.** HEK cells were co-transfected with p62-Myc and HA-IκBα wildtype (WT), or HA-IκBα with all the 9 Ks mutated to Rs (9KR), or mutated with just K238 to R (K238R). Cells were then left untreated (Ctrl) or treated with NEM (10 μM) for 5 min before harvesting. Cells were then harvested in RIPA buffer, and then subjected to IP with HA-agarose followed by immunoblotting. **D**. HEK cells overexpressing HA-IκBα and p62-Myc were either untreated or treated with NEM followed by in-cell crosslinking with 5 mM DSS or 0.05 mM SIAB, lysed in RIPA, then subjected to IP and immunoblotting. White arrows indicate IκBα monomers that disappear upon DSS-crosslinking with NEM treatment. **E.** HEK cells overexpressing HA-IκBα and p62-Myc were treated with NEM followed by in-cell crosslinking with indicated concentrations of DSS, lysed in RIPA, and then subject to IP and immunoblotting.

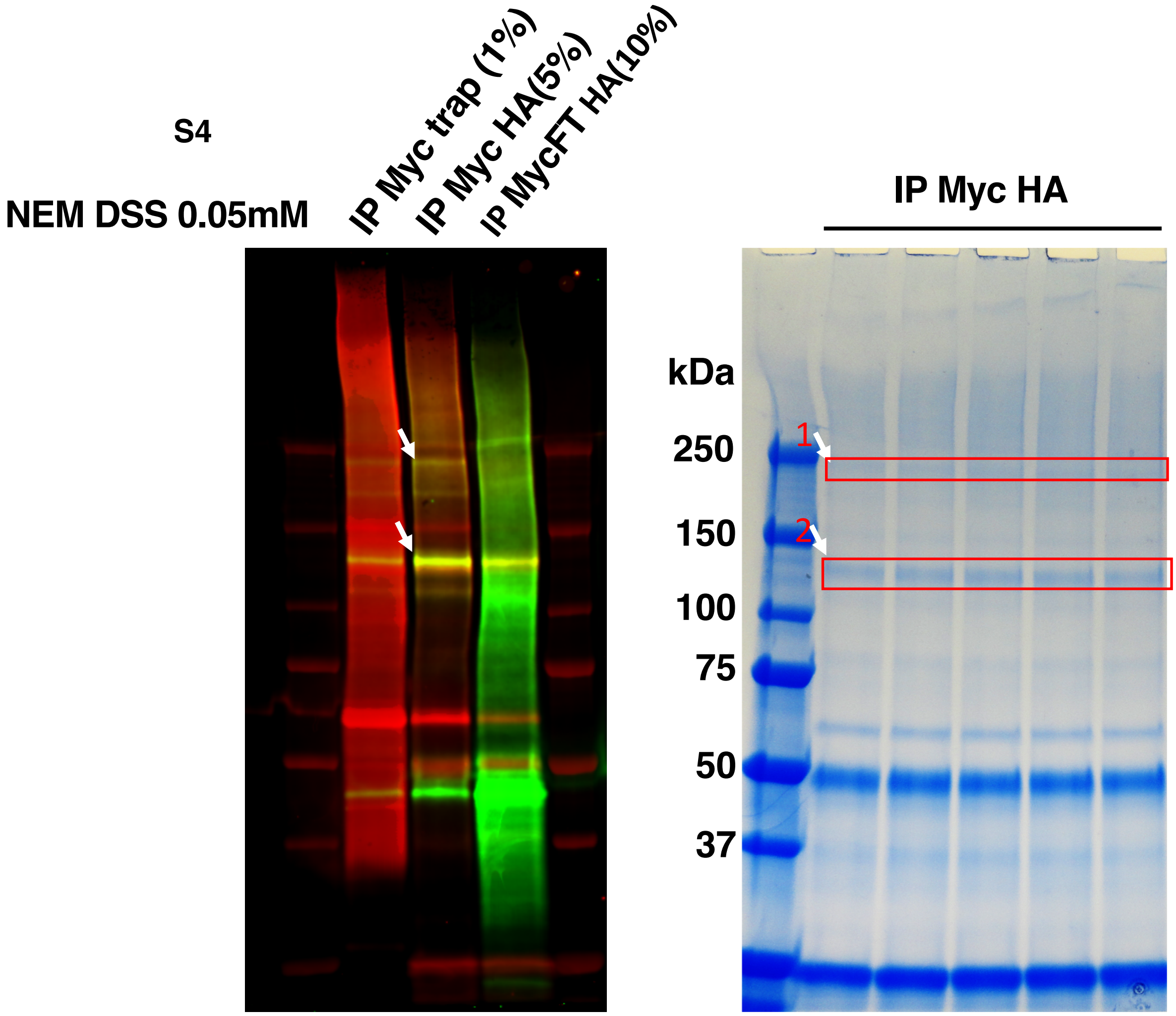


***FIGURE S4. Gel images showing bands used for DSS X-linking LC-MS/MS.*** HEK cells expressing HA-IκBα and p62-Myc were first treated with NEM, then in-cell DSS (0.05mM) X-linking for 30 min. Ten 60mm plates were pooled, lysed in RIPA buffer. ~ 50mg lysates were used for tandem Myctrap (500 μl beads) and HA-agarose (100 μl agarose) IP. A small fraction of eluates and flowthrough (FT) were used for immunoblotting. Final eluates were run across 5 lanes on SDS-PAGE. Indicated bands were excised and in-gel digested with trypsin + Lys-C, and the extracts subjected to LC-MS/MS to detect Xlinked peptides.

S5

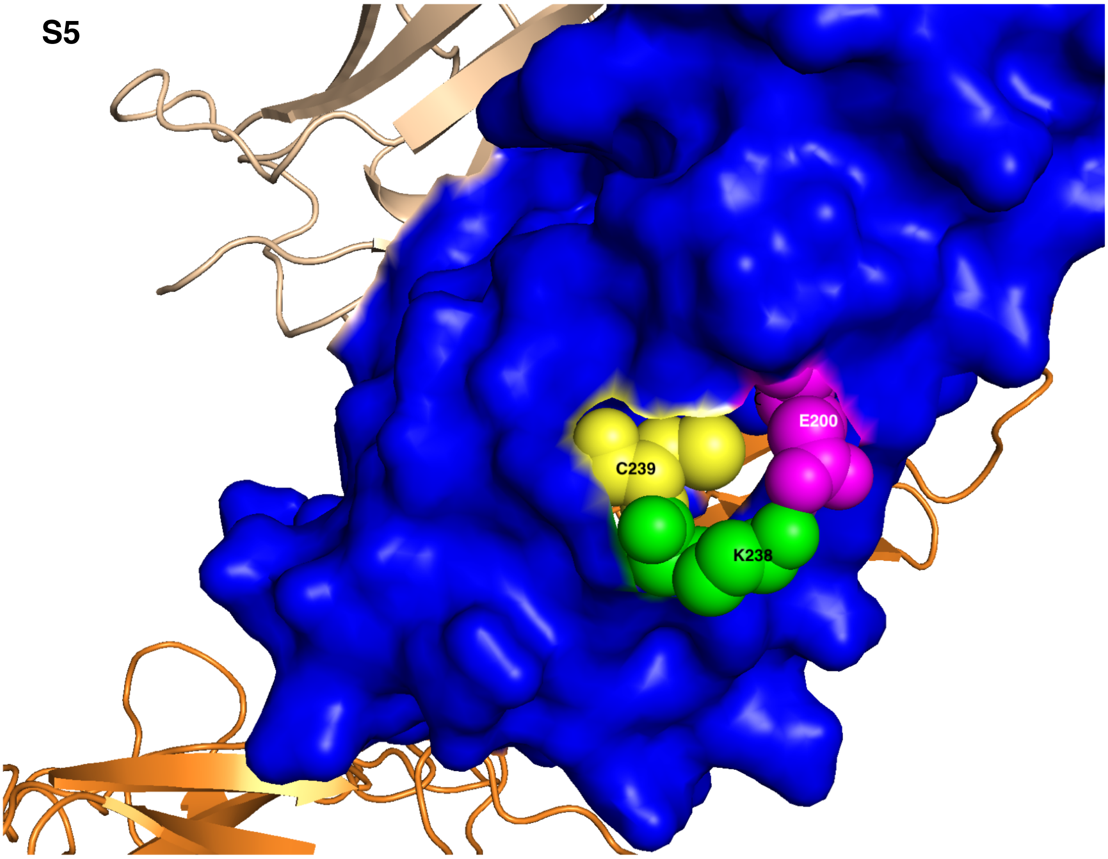


***FIGURE S5.*** Close-up of Fig 5A showing residues K238, C239 and E200.

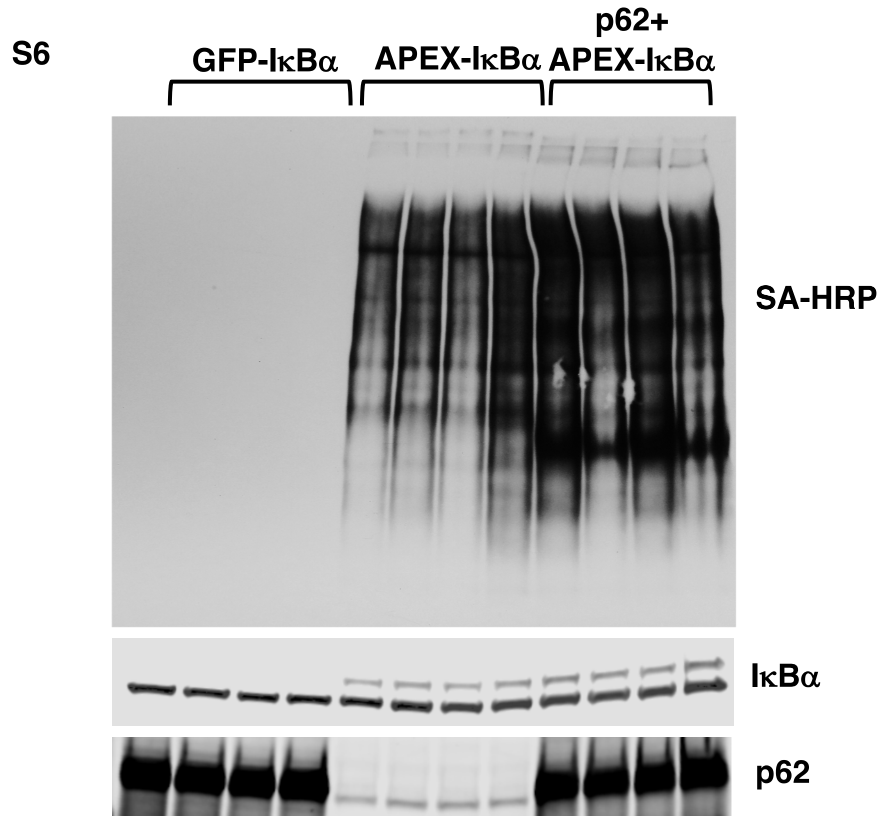


***FIGURE S6. APEX-mediated proximity biotinylation-labeling analyses:*** HEK cells expressing GFP-IκBα with p62-Myc, APEX-IκBα alone, or APE-IκBα with p62-Myc were treated with $H_2O_2$ and Biotin-phenol to initiate APEX-mediated proximity biotinylation labeling. Four biological replicates were performed for each condition. Cells were lysed in RIPA and subject to immunoblotting to examine biotinylation before proceeding to enrichment with SA-beads.

**S7A** HepG2

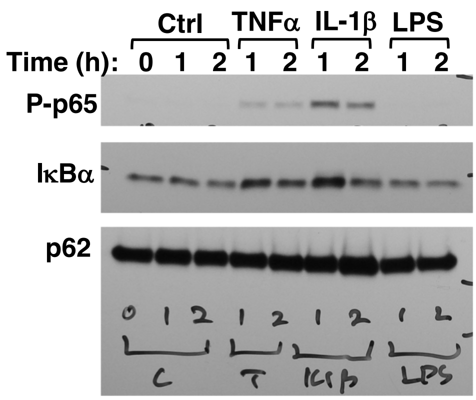


**S7B**

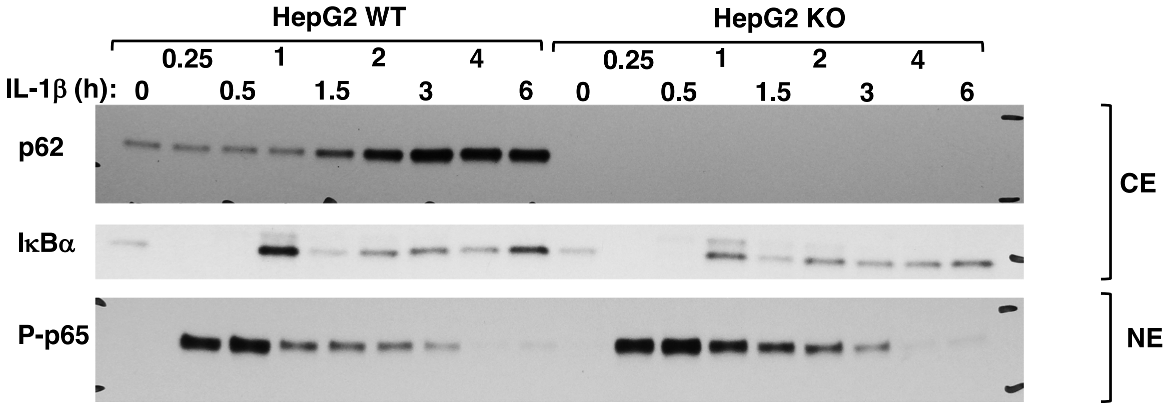


***FIGURE S7. HepG2 cells exhibit a strong NF-κB activation upon IL-1β stimulation***. **A**. HepG2 cells were first rested in no FBS medium overnight, then treated with TNFα or IL-1β or LPS at the indicated times. Cell lysates were subjected to immunoblotting analyses. **B**. WT or p62 CRISPR knockout (KO) HepG2 cells were first rested in no FBS medium overnight, then treated with IL-1β at the indicated times. Cytosol extracts (CE) and nuclear extracts (NE) were subjected to immunoblotting analyses.

## S8

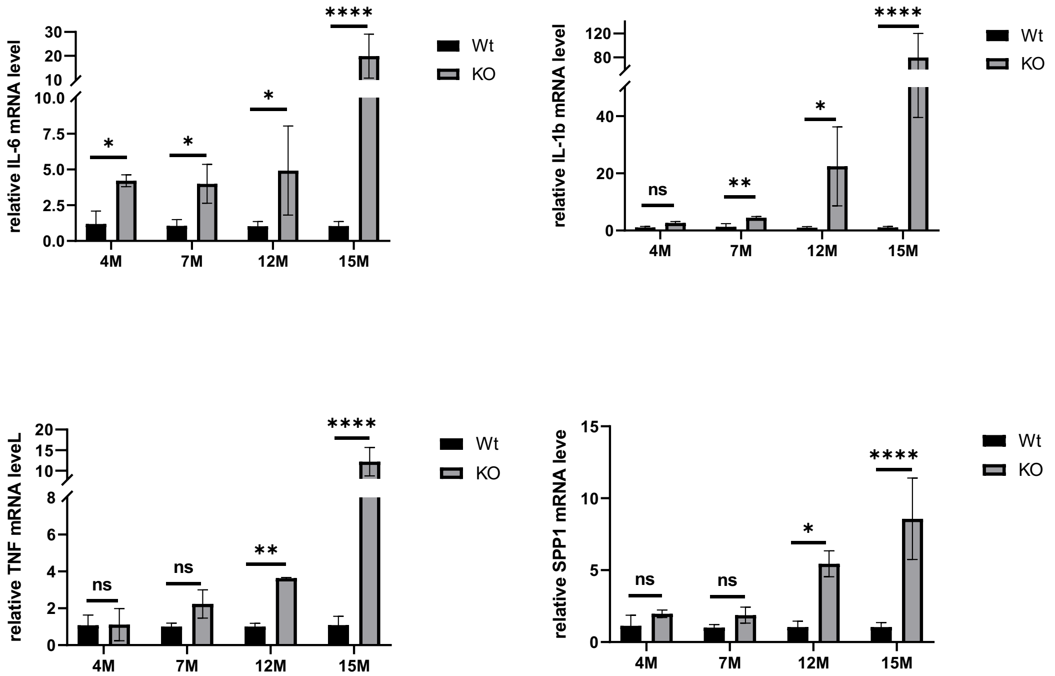


***FIGURE S8. Relatively enhanced liver inflammatory markers [IL-6, IL-1$\beta$, tumor necrosis factor-$\alpha$ (TNF-$\alpha$), and SPP1] in WT and p62mut-mice upon aging.*** Total RNA (2.5 μg) from age-matched mouse livers was extracted as described (17) and subjected to reverse transcription based on equal amounts of RNA using the SuperScript$^{TM}$ IV VILO$^{TM}$ Master Mix with ezDNase kit (Invitrogen, REF #11766050). Quantitative PCR (qRT-PCR) analyses were then performed in triplicate for each sample using the PowerUp$^{TM}$ SYBR$^{TM}$ Green Master Mix (Thermo Fisher Scientific, Cat #A25742) according to the manufacturer's instructions with the primers summarized in **Table S1**. The relative expression of genes of interest was compared with that of the reference gene, glyceraldehyde-3-phosphate dehydrogenase (GAPDH). Data were analyzed using GraphPad Prism software, and the statistical significance determined by the Student's t-test vs age-matched WT, as follows: NS, not significant; *p < 0.05, **p < 0.01. ****p < 0.0001.

**Table S1. List of All Primers Used for Real-Time PCR (Fig. S8)**

| no | Gene name | Sequence (5' to 3') |
|---|---|---|
| 1 | qTNF-F | AGG GTC TGG GCC ATA GAA CT |
| 2 | qTNF-R | CCA CCA CGC TCT TCT GTC TAC |
| 3 | qIL6-F | CTC TGC AAG AGA CTT CCA TCC AGT |
| 4 | qIL6-R | GAA GTA GGG AAG GCC GTG G |
| 5 | qIL1B-F | TCT TTG AAG TTG ACG GAC CC |
| 6 | qIL1B-R | TGA GTG ATA CTG CCT GCC TG |
| 7 | SPP1-F | ATT CTG GCA GCT CAG AGG AG |
| 8 | SPP1-R | CTG TGG CGC AAG GAG ATT |
| 9 | qGAPDH-F | TGT GTC CGT CGT GGA TCT GA |
| 10 | qGAPDH-R | TTG CTG TTG AAG TCG CAG GAG |

**S9.**

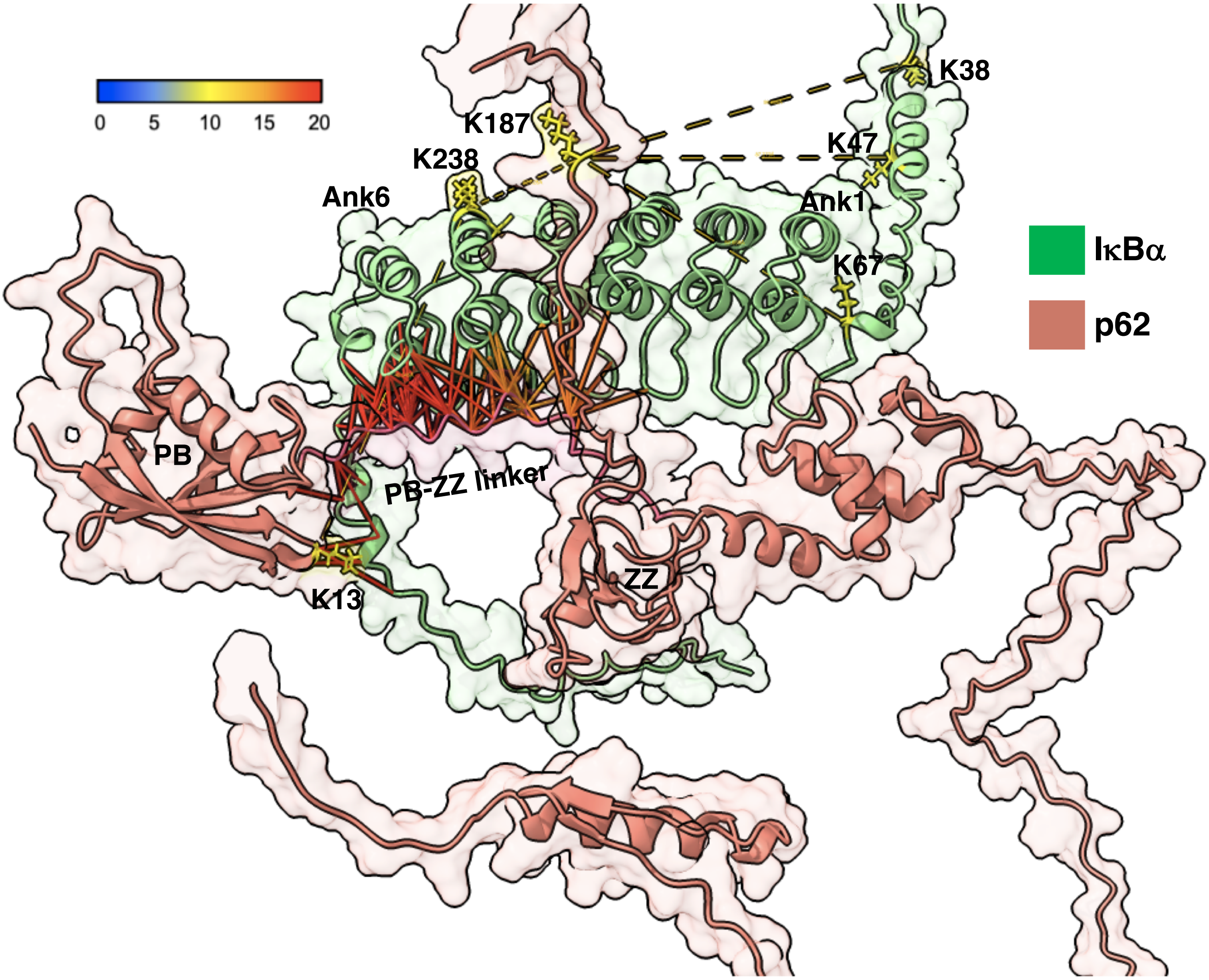


***FIGURE S9. AlphaFold of the IκBα-p62 complex depicting the Predicted Aligned Errors (PAE) values.*** Other details as in Fig.10.
